# Phenotypic screening of the ReFrame Drug Repurposing Library to discover new drugs for treating sickle cell disease

**DOI:** 10.1101/2022.06.23.497377

**Authors:** Belhu Metaferia, Troy Cellmer, Emily B. Dunkelberger, Quan Li, Eric R. Henry, James Hofrichter, Dwayne Staton, Matthew M. Hsieh, Anna K. Conrey, John F. Tisdale, Arnab K. Chatterjee, Swee Lay Thein, William A. Eaton

**Affiliations:** Laboratory of Chemical Physics, National Institute of Diabetes and Digestive and Kidney Diseases; Office of the Clinical Director, National Institute of Diabetes and Digestive and Kidney Diseases; Molecular and Clinical Hematology Branch, National Heart, Lung, and Blood Institute; National Institutes of Health; Sickle Cell Branch, National Heart, Lung, and Blood Institute; National Institutes of Health, Bethesda, MD, 20892; Calibr at Scripps Research, La Jolla, CA 92037 USA

**Keywords:** sickle cell disease, anti-sickling drugs, hemoglobin S, phenotypic drug discovery, high throughput drug screening

## Abstract

Stem-cell transplantation and genetic therapies offer potential cures for patients with sickle cell disease (SCD) but these options require advanced medical facilities and are expensive. Consequently, these treatments will not be available to the vast majority of patients suffering from this disease for many years. What is urgently needed now is an inexpensive oral drug in addition to hydroxyurea, the only successful drug approved by the FDA that inhibits sickle-hemoglobin polymerization. Here we report results of the first phase of our phenotypic screen of the 12,657 compounds of the Scripps ReFrame drug repurposing library using a recently developed high-throughput assay to measure sickling times following deoxygenation to 0% oxygen of red cells from sickle trait individuals. The ReFrame library is a very important collection because the compounds are either FDA-approved drugs or have been tested in clinical trials. From dose-response measurements, 106 of the 12,657 compounds exhibit statistically significant anti-sickling at concentrations ranging from 31 nM to 10 μM. Compounds that inhibit sickling of trait cells are also effective with SCD cells. As many as 21 of the 106 anti-sickling compounds emerge as potential drugs. This estimate is based on a comparison of inhibitory concentrations with free concentrations of oral drugs in human serum. Moreover, the expected therapeutic effect for each level of inhibition can be predicted from measurements of sickling times for cells from individuals with sickle-syndromes of varying severity. Our results should motivate others to develop one or more of these 106 compounds into drugs for treating SCD.

**Significance Statement:** The vast majority of patients suffering from sickle cell disease live in under-resourced countries. Consequently, advanced medical facilities required for curative therapies, such as stem cell transplantation and gene therapy, will be unavailable to them for a long time. Hydroxyurea, approved by the FDA in 1998, is the only effective drug that inhibits polymerization of the mutant hemoglobin S that stiffens and distorts (“sickles”) red cells, the root cause of the pathology. What is urgently needed now for these patients are additional, inexpensive oral anti-sickling drugs. Our high throughput phenotypic screen of the ReFrame drug repurposing library reported here discovered 106 compounds that are anti-sickling. On a statistical concentration basis, as many as 21 are predicted to be potential drugs.

---

An abnormal hemoglobin (HbS) causes sickle cell disease (SCD), the first example of a “molecular disease” reported by the legendary chemist, Linus Pauling, in 1949(1, 2). Pauling’s discovery gave birth to the field of molecular medicine. This mono-genetic disease results from a single nucleotide change from A to T in the codon (GAG) of the gene for the hemoglobin beta chain. The mutation results in the replacement of a negatively charged glutamate with a neutral, hydrophobic valine on the molecular surface, which creates a sticky patch that promotes polymerization of HbS upon deoxygenation to form fibers (3, 4). HbS polymerization in the tissues is the root cause of SCD pathology because the fibers stiffen and distort (“sickle”) red cells. Sickled cells can cause vaso-occlusion in the smallest vessels of almost every tissue, resulting in both acute and chronic organ damage and episodes of pain that are so severe that they are called a sickle cell crisis(5, 6). In addition, the large and often irreversible structural changes in the red cell cytoskeleton associated with sickling in the tissues and unsickling in the lungs causes red cell fragility and consequent hemolysis that underlies the chronic hemolytic anemia. Inhibition of HbS polymerization is therefore the most rational drug treatment for the disease(7).

Hydroxyurea, approved by the FDA in 1998, is currently the only successful anti-sickling drug. Although very beneficial, it is not curative and very much underutilized in low resource settings where the incidence of SCD is high, such as sub-Saharan Africa(8). Hydroxyurea is effective because it induces the synthesis of fetal hemoglobin, which inhibits polymerization (7, 31), but it is not induced in all cells. Thus, additional inexpensive oral drugs that inhibit sickling in all cells are urgently needed because the vast majority of patients in the world do not have and for many years will not have the necessary advanced medical facilities and resources required for curative therapies such as hematopoietic stem cell transplantation or gene therapy(9, 10). Now that large compound libraries are available, basic research scientists can engage in drug development using screening assays that are patho-physiologically relevant. We have recently developed a relevant high throughput assay(11) to screen the Scripps California Institute of Biomedical Research (Calibr) ReFrame drug repurposing library. This library is a unique and very important collection because the compounds are either FDA-approved drugs for other indications or have been tested in clinical trials with extensive drug annotation in an open-access format (reframedb.org)(12). Consequently, compounds that inhibit sickling at serum concentrations found in humans, without known side effects that would be deleterious for SCD patients, can be rapidly brought to clinical trials without extensive pre-clinical studies that markedly slow drug development. Since there is no target, this approach is referred to as phenotypic drug discovery (13). In this work, we present our results so far from screening the 12,657 compounds of the Calibr library using our sickling assay(11).

Our previous screening assay used a 96-well plate format and laser photodissociation of carbon monoxide (CO) saturated sickle trait cells to rapidly create deoxyHbS in less than one second (14). The assay is extremely sensitive because it determines the delay time prior to sickle fiber formation for each cell, but it is low throughput and requires the use of sodium dithionite, a very strong reducing agent that could react with the test compound(14, 15). We were therefore motivated to develop a high throughput assay under more physiological conditions. Our current assay uses a 384 well plate format and is based on deoxygenation of sickle trait cells with nitrogen to 0% oxygen and acquisition of images of 100-300 cells every 15 minutes using an Agilent “Lionheart” automated microscope system(11). With one instrument, up to ∼1300 compounds at a single concentration can be tested in one week, making it relatively high throughput for drug screening; we now have 4 such instruments. As in the previous CO photolysis assay, the time at which individual cells change shape (sickle) is determined using our automated image algorithms for characterizing red cell morphology, refined by machine learning(11).

There are many advantages to using cells from donors with sickle trait, the heterozygous condition(14). The delay time for polymerization is about 1000-fold longer for trait cells than for SCD cells because of the much lower fraction of HbS in trait cells (35-40% HbS in trait compared to 85-95% in SCD)(16, 17). With longer delay times, there are many fewer primary nucleation events, resulting in only a few domains of fibers (18, 19). Consequently, cellular distortion is much greater (20), making the determination of sickling times more accurate. There are several additional advantages in using trait cells, including (i) the fact that they are less damaged and therefore are a much more homogeneous cell population than SCD cells(6, 21) and (ii) that there are many more blood donors, who very often have relatives with SCD and are therefore willing to donate as frequently as requested. While polymerization in trait cells is much slower, there is no evidence that the mechanism of sickle fiber formation is different(19). Nevertheless, it is important that we demonstrate that inhibitors of trait cell sickling also inhibit sickling of SCD cells.

There is every reason to expect that new drugs will be discovered from our screen because it tests for all but one of the five independent approaches that have been proposed to inhibit sickling as therapy(7). These four classes of inhibitory mechanisms, discussed in detail in reference 7, are: (i) blocking inter-molecular contacts in the sickle fiber, (ii) destabilizing the fiber by decreasing 2,3-diphosphoglycerate (2,3-DPG)(22, 23), with added destabilization resulting from the concomitant increase in intracellular pH(24–26), (iii) decreasing the intracellular hemoglobin concentration, for example by swelling red cells(14, 27), and (iv) preferentially shifting the R-T quaternary equilibrium toward the non-polymerizing R conformation with a drug that both binds and dissociates very rapidly from the R quaternary conformation (15, 19, 28, 29). Since the assay measures inhibition of mature cells in peripheral blood, it does not test for compounds that induce fetal hemoglobin (HbF) synthesis in stem cells (30–35). In order to assess this 5^th^ independent approach, we make considerable use of the assay to measure sickling of blood from individuals with SCD and sickle trait on various protocols and from patients who have been treated by using a virus to add globin genes to their stem cells for the non-polymerizing HbF mimicking mutant, HbA(T87Q)(36).

## Results

### Images of sickle trait cells upon deoxygenation

Figure 1 shows 3 typical images at 13 hrs after the start of deoxygenation, one for the negative control with 85% of cells sickled, one for the positive control with no cells sickled, and one for a test compound that shows 40% of cells sickled. To have a sufficient number of cells in each image at acceptable resolution for the automated image analysis, a 20x objective with a numerical aperture of 0.45 was employed. Consequently, the resolution of the images is sub-optimal. Nevertheless, the cellular distortions are more than sufficient to determine at each time point whether a cell is sufficiently distorted to label it as sickled. The three most obvious changes are the loss of a circularity, the loss of a more transparent center because of the loss of concavity in the middle of the cell, and the decrease in the projected area of the cell. The time at which a cell sickles is determined by a change in at least two of these metrics compared to the previous time point. The output of the analysis is the fraction of cells sickled as a function of time. In the supplementary information (S.I.) we show a video of images following the start of deoxygenation,

**Figure 1.**
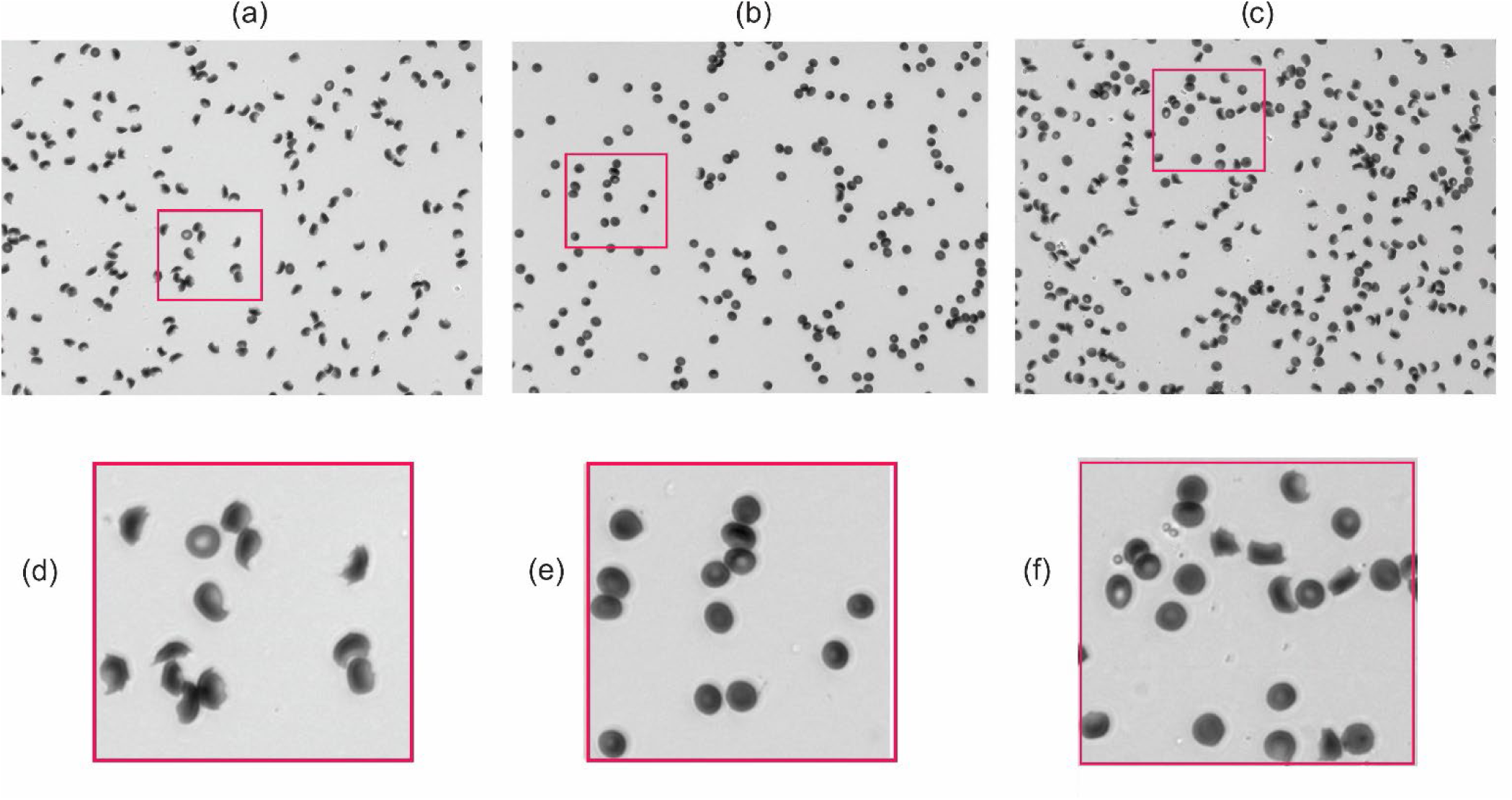
Images of sickle trait cells. (two column final figure). (a) Negative control at end of experiment (13 hrs after start of deoxygenation). (b) Positive control at end of experiment (13 hrs after start of deoxygenation). (c) Well containing 10 μM of phanquinone (13 hrs after start of deoxygenation). The fraction sickled in the negative control for this well is 0.85, while the fraction in the well containing phanquinone is 0.40. There are 172, 175, and 236 cells in each of these images, respectively. (d,e,f) Zoomed in sections of images in panels a,b,c, respectively.

### Fraction sickled vs time and dose-response results

Figure 2a shows the time courses for different concentrations of the test compound, calcymycin, as well as for the negative and positive controls, of the fraction sickled following the start of deoxygenation to 0% oxygen with nitrogen to induce HbS polymerization and sickling. Also shown with these sickling curves are the singular value decomposition (SVD) of the curves and the dose-response results (panel b), which showed 50 percent inhibition (IC50) at a concentration of 2.1 ± 0.1 μM. SVD is a powerful matrix-analytic method for averaging data to eliminate noise and provides the best least-squares representation of the results with smooth curves that simplifies the quantitative analysis of massive data sets (37). SVD of the measured sickling curves is described in detail in the **Methods** section. The three colors correspond to the results for red cells from three different sickle trait donors. This set of experiments is not representative but shows how good the data can get when almost every aspect of the experiment and image analysis works nearly perfectly. In this case the differences among the negative controls are much larger than usual because of large differences in hemoglobin composition and intracellular hemoglobin concentration (MCHC), which are the two most important factors that determine the rate and extent of sickling. (MCHC is the abbreviation for the clinical laboratory term, mean corpuscular hemoglobin concentration). Cells from the green donor sickle the most because they contain the largest percentage of HbS and the highest MCHC. Nevertheless, the fraction sickled as a function of compound concentrations relative to the negative control is very similar for all three donors (Fig. 2b).

**Figure 2.**
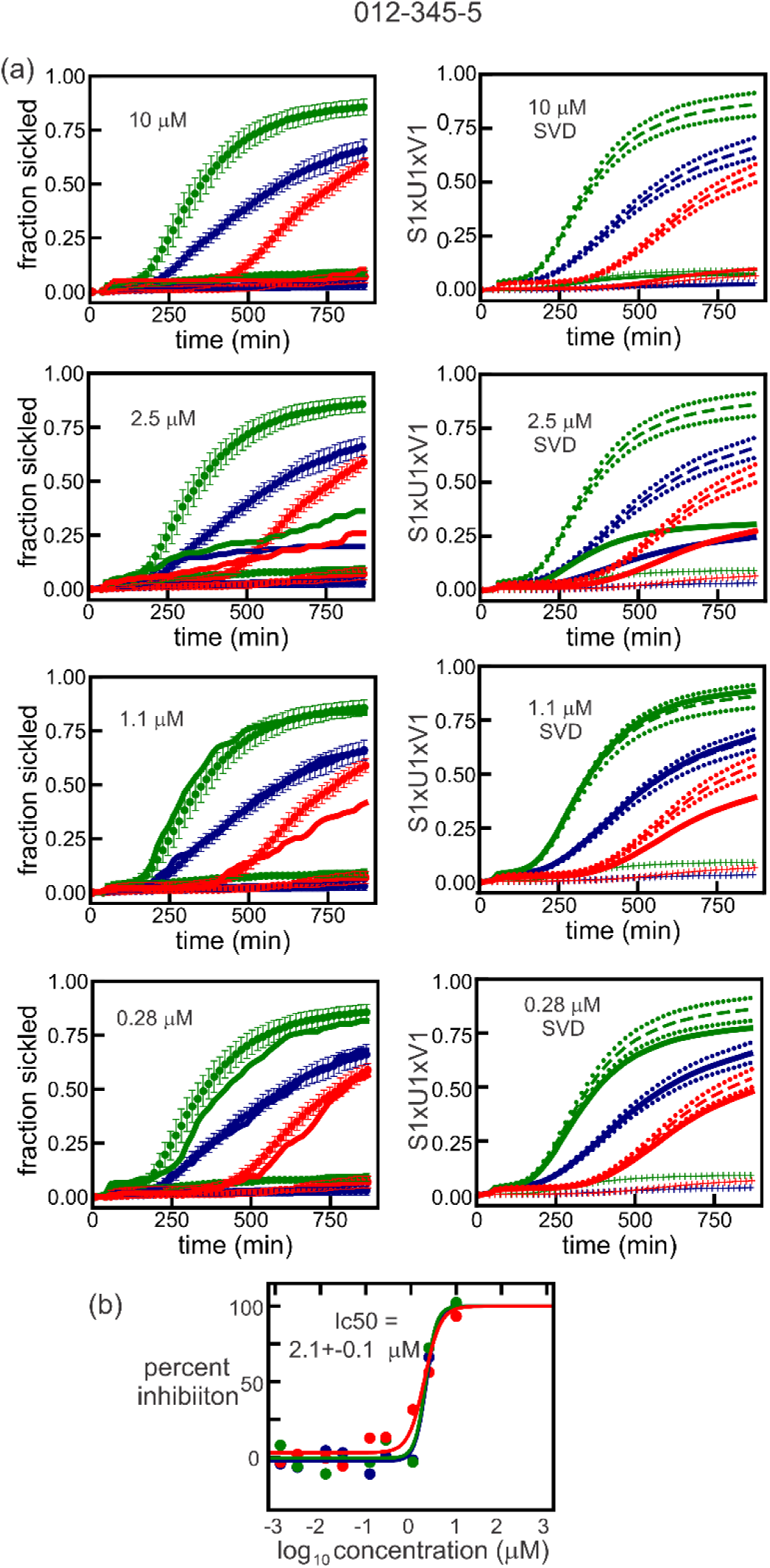
Non-representative dose response data for ReFrame compound calcimycin, an anti-biotic. (one column figure) (a) Time course of the fraction sickled following the start of deoxygenation with nitrogen to induce HbS polymerization and sickling. Sickling curves at four different concentrations of calcimycin. Panels on the left are the measured sickling curves, while panels on the right show the curves represented by the first component of the SVD. The continuous curves in both left and right panels are the sickling curves for the single wells containing the test compound. The three colors represent samples from three different sickle trait donors with hemoglobin compositions as percentage of total hemoglobin: green donor: HbA 56.0, HbS 38.0, HbA2 4.0, HbF 1.2, MCHC 34.2 g/dL; blue donor: HbA 62.7, HbS 33.7, HbA2 3.5, HbF<1.0, MCHC 28.6 g/dL; red donor: HbA 59.7, HbS 37.8, HbA2 3.8, HbF<1.0, MCHC 26.6 g/dL. In the left panels, the upper points are the average and standard deviations from the average for 12 negative controls, while the lower points are the average and standard deviations from the average of 6 positive controls. In the right panels, the long-dashed curves are the average sickling curves (S1xU1xV1) of the negative controls with standard deviations indicated by the short-dashed curves. The lower points and vertical bars are the average and standard deviations for the positive controls. (b) Points are measured percent inhibition defined by equation (1) for 3 different donors, while the continuous curves are the fits to the data using equation (2) (see **Methods** section).

Figure 3 is more representative of the results for all the compounds measured. Four additional representative data sets are given in the S.I. All the dose response plots, as in panels (b) of Figures 2 and 3, for the 99 of the 106 compounds found to be anti-sickling are given in the S.I.

**Figure 3.**
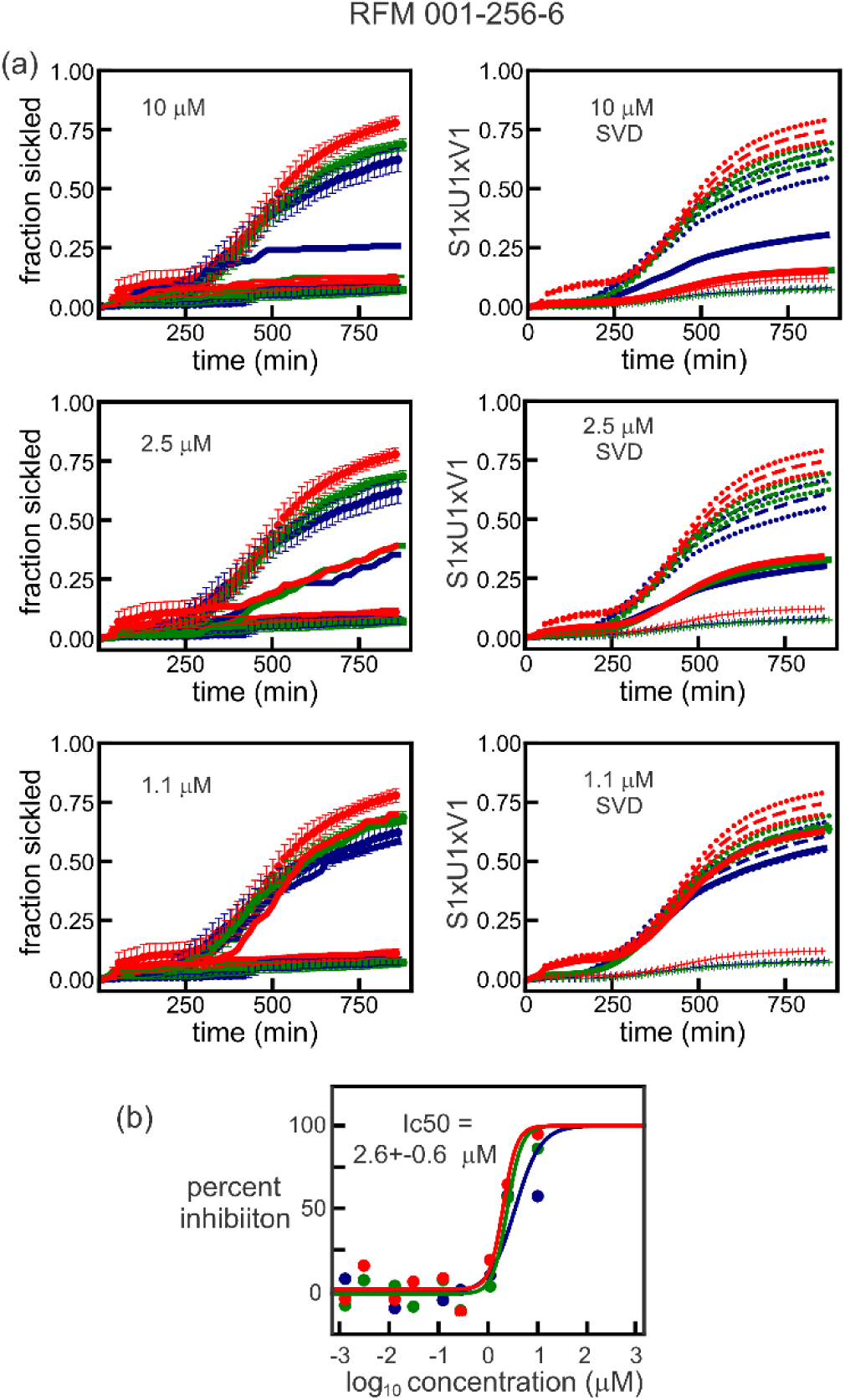
More representative dose response data for ReFrame compound NSC-150117, phosphatase inhibitor. (one column final figure). The three colors represent samples from three different sickle trait donors with hemoglobin compositions: blue donor HbA 59.7, HbS 37.8, HbA2 3.8, HbF<1.0, MCHC 26.6 g/dL; green donor: HbA 59.8, HbS 35.5, HbA2 3.8, HbF < 1.0, MCHC 33.6 g/dL; red donor: HbA 56.1, HbS 40.0, HbA2 3.7,HbF <1.0, MCHC 29.8 g/dL. Remainder of legend is same as for Figure 2.

Table 1 summarizes our main results. It gives the compound name, the concentration that produces 50% inhibition (IC50), and the lowest concentration that produces statistically significant inhibition (LIC), defined by 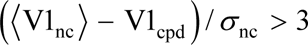, where 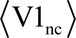 is the average value of the amplitude (V1) of U1 for the sickling curves of the negative controls (nc), V1_cpd_ is the amplitude of U1 for the sickling curve of the test compound, and σ_nc_ is the standard deviation of 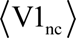. See **Methods** for description of the U1 and V1’s, the first components of the **U** and **V** matrices of the singular value decomposition.

**Table 1.** List of anti-sickling compounds with lowest inhibitory concentrations (LIC) less than 10 μM.

| RFM number | Name of Compound | IC<sub>50</sub> (μM)<sup>a</sup> | LIC (μM)<sup>b</sup> |
| --- | --- | --- | --- |
| RFM-000-126-3 | Ammonium Trichlorotellurate <sup>c,d</sup> | 0.032 ± 0.015 | 0.031 |
| RFM-010-363-9 | Niferidil <sup>c,h</sup> | 10.5 | 0.031 |
| RFM-004-015-3 | Mavatrep | 0.23 ± 0.14 | 0.123 |
| RFM-006-618-2 | Manoalide <sup>c</sup> | 0.33 ± 0.09 | 0.278 |
| RFM-007-990-3 | HR325 <sup>c</sup> | 3.0 ± 0.86 | 0.278 |
| RFM-009-396-9 | NSD-721 <sup>c</sup> | 13 ± 2.0 | 0.278 |
| RFM-010-342-4 | FT-11 <sup>c</sup> | 1.3 ± 1.0 | 0.278 |
| RFM-014-044-3 | Mitapivat <sup>c</sup> | 0.25 ± 0.16 | 0.278 |
| RFM-001-577-0 | ARA DMAP <sup>c</sup> | 3.9 ± 1.1 | 1.1 |
| RFM-002-851-3 | Pafuramidine <sup>c</sup> | 6.0 ± 5.8 | 1.1 |
| RFM-002-987-8 | Cisplatin <sup>c,d,e</sup> | 1.3 ± 0.2 | 1.1 |
| RFM-004-808-8 | Cloturin <sup>c,g</sup> | 9.7 ± 0.5 | 1.1 |
| RFM-004-814-6 | EF-4040 <sup>c</sup> | 1.7 ± 0.4 | 1.1 |
| RFM-005-694-0 | Cyclovalone <sup>c</sup> | 1.7 ± 0.7 | 1.1 |
| RFM-007-932-3 | VML 530 | 1.5 ± 0.2 | 1.1 |
| RFM-008-390-9 | Peruvoside <sup>c</sup> | NA<sup>f</sup> | 1.1 |
| RFM-009-283-1 | ONO-7746 | 3.8 ± 2.0 | 1.1 |
| RFM-010-785-7 | BMS-919373 <sup>d,e,g</sup> | 6.4 ± 8.1 | 1.1 |
| RFM-011-401-2 | KUX-1151 <sup>c</sup> | 5.3 ± 5.1 | 1.1 |
| RFM-011-459-0 | NT-113 | 3.6 ± 2.8 | 1.1 |
| RFM-012-385-3 | Curcumin | 0.50 ± 0.14 | 1.1 |
| RFM-012-490-3 | AEG-33783 <sup>c,d</sup> | 1.4 ± 0.5 | 1.1 |
| RFM-000-489-7 | VRT-532 <sup>c</sup> | 5.8 ± 0.7 | 2.5 |
| RFM-000-943-8 | Licochalcone A | 4.3 ± 2.4 | 2.5 |
| RFM-001-256-6 | NSC-150117 | 2.6 ± 0.7 | 2.5 |
| RFM-002-051-9 | AT-101 <sup>d</sup> | 2.4 ± 0.1 | 2.5 |
| RFM-002-220-8 | Darapladib <sup>d,e</sup> | 2.0 ± 0.4 | 2.5 |
| RFM-002-539-8 | Oxyfedrine Hydrochloride | 2.9 ± 1.1 | 2.5 |
| RFM-004-129-2 | Octenidine <sup>c,d,e</sup> | NA<sup>f</sup> | 2.5 |
| RFM-004-629-7 | WP-1066 <sup>c</sup> | 3.9 ± 1.2 | 2.5 |
| RFM-004-737-0 | Butein | 10 ± 3 | 2.5 |
| RFM-005-775-0 | Etacrynic Acid <sup>c</sup> | 3.3 ± 0.9 | 2.5 |
| RFM-006-284-0 | TS-80 <sup>c</sup> | 3.1 ± 1.6 | 2.5 |
| RFM-007-863-7 | SC-1 | 10 ± 5 | 2.5 |
| RFM-007-872-8 | Tucaresol | 3.1 ± 0.3 | 2.5 |
| RFM-008-587-0 | CBS-1114 | 7.2 ± 5.2 | 2.5 |
| RFM-009-852-2 | YM 57158 | 2.5 ± 0.3 | 2.5 |
| RFM-010-353-7 | Allisartan <sup>c</sup> | 7.8 ± 4.6 | 2.5 |
| RFM-010-371-9 | Elisartan Potassium | 5.6 ± 4.0 | 2.5 |
| RFM-010-692-3 | Samixogrel | 4.2 ± 2.0 | 2.5 |
| RFM-011-023-6 | AM-103 <sup>c,d,g</sup> | 4.5 ± 2.6 | 2.5 |
| RFM-011-462-5 | ODM-103 <sup>c,g</sup> | 9.0 ± 2.2 | 2.5 |
| RFM-011-465-8 | ON-09310 | 5.9 ± 1.7 | 2.5 |
| RFM-011-681-4 | P-7435 <sup>c</sup> | NA<sup>f</sup> | 2.5 |
| RFM-011-880-9 | AZD-4769 | 2.5 ± 0.9 | 2.5 |
| RFM-012-345-5 | Calcimycin | 2.1 ± 0.1 | 2.5 |
| RFM-012-436-7 | Vadadustat <sup>c</sup> | 7.9 ± 6.3 | 2.5 |
| RFM-012-604-5 | Metabromsalan <sup>c</sup> | 8.0 ± 2.5 | 2.5 |
| RFM-012-837-0 | IMD-0560 | 2.3 ± 1.1 | 2.5 |
| RFM-012-931-7 | laflunimus | 7.2 ± 3.7 | 2.5 |

<sup>a</sup>concentration that produces 50% inhibition
<sup>b</sup>lowest concentration of statistically significant inhibition
<sup>c</sup>LIC is anti-sickling for cells from 2 of 3 donors
<sup>d</sup>Compound causes lysis at high concentration, anti-sickling at lower concentration
<sup>e</sup>Incubated for 8 hours
<sup>f</sup>IC50s could not be determined for these 3 compounds because of an insufficient number of data points higher than 0 % inhibition
<sup>g</sup>IC50s could only be determined for 2 of the 3 donor samples for these compounds because of an insufficient number of data points higher than 0 % inhibition
<sup>h</sup>IC50s could only be determined for 1 of the donor samples for these compounds because of an insufficient number of data points higher than 0 % inhibition
<sup>i</sup>See (15, 29)

### Comparing sickling inhibition of sickle trait and SCD cells

To determine whether compounds that inhibit sickling of cells from sickle trait donors also inhibit sickling of cells from SCD patients, a comparison was carried out for several of the anti-sickling compounds discovered in the screen. Compounds for these experiments were purchased from commercial sources (see company names in **Materials and Methods**). In order to have roughly comparable sickling curves for the negative controls of both SCD and trait cells, SCD cells were deoxygenated to 5% oxygen to slow polymerization; trait cells were deoxygenated to 0% oxygen as in the screen. Figure 4 shows a comparison of the sickling curves for trait and SCD cells for 14 of the anti-sickling compounds in Tables 1 and S1.

**Figure 4.**
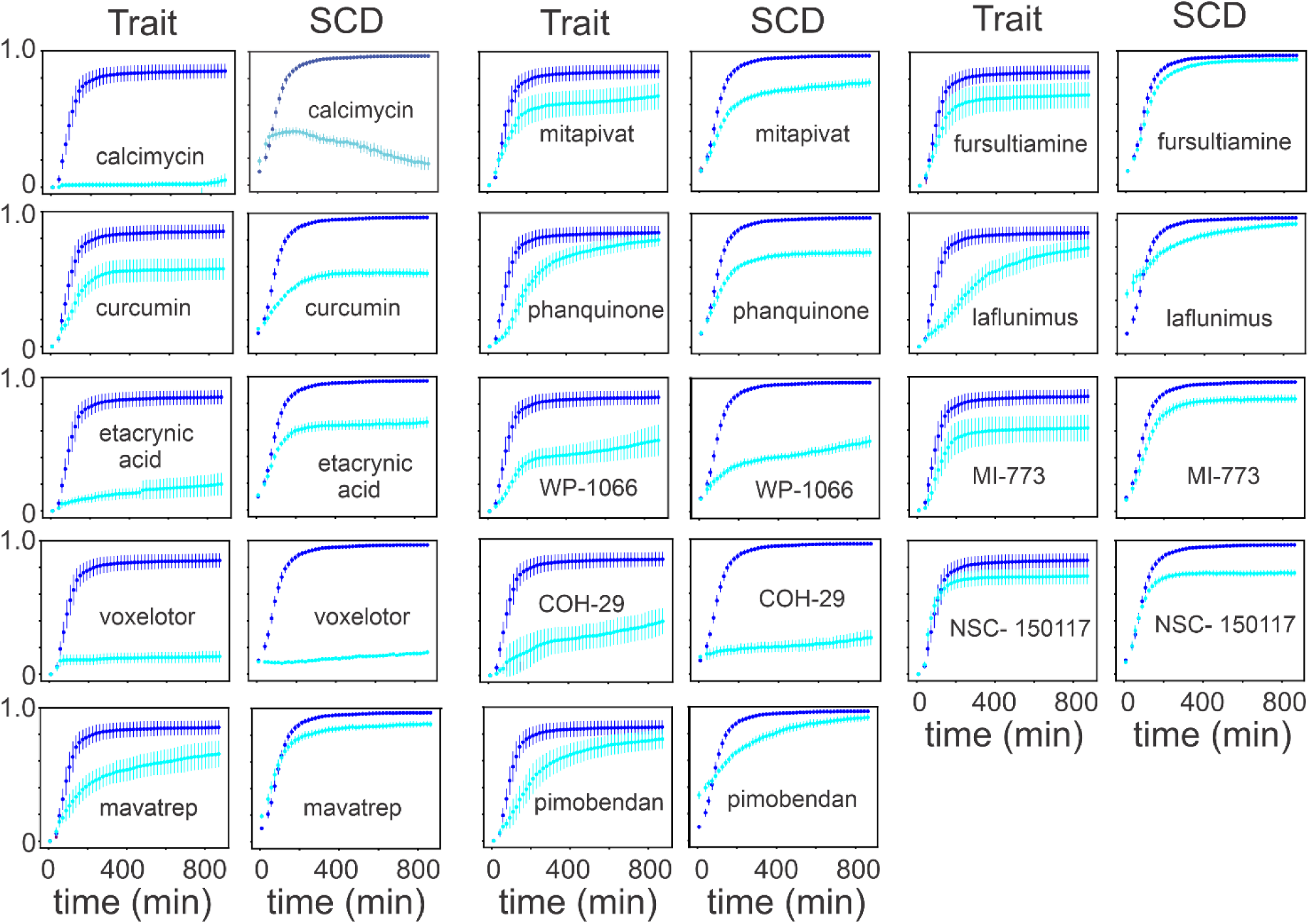
Comparing anti-sickling compounds in sickle trait and SCD cells. (two column l figure. Fraction sickled vs time following deoxygenation to 0% oxygen for cells from sickle trait individual (HbA 56.1%, HbS 40%, HbA2 3.7%, HbF <1.0%, MCHC 29.5 g/dL) and deoxygenation to 5% oxygen for cells from for SCD patient (HbS 91 %, HbA2 4.2 %, HbF 4.8 %, MCHC 34.8 g/dL) incubated for 3 hrs at a concentration of 10 μM for 14 of the 106 anti-sickling compounds listed in Table 1. In all the plots the *x* axis runs from 0 to 900 min. and the *y* axis from 0 to 1, both with slight offsets from 0. The upper curves (blue) are negative controls; lower curves (cyan) are for test compounds. The standard deviations are larger for trait cells because the average number of cells in the images was about 2-fold less than for SCD cells.

### Comparing anti-sickling concentrations with serum concentrations of oral drugs in PDR

The distribution of maximum ***free*** concentrations (C_max_) for oral drugs in the Physician’s Desk Reference (PDR) is shown in Figure 5a. The free C_max_ is the maximum free concentration of a drug for the given dosage regime. Figure 5b shows the number of anti-sickling compounds at each concentration for which there is statistically significant inhibition of sickling, as well as the probable number in parentheses that would be expected to be oral drugs based solely on the concentration distribution in Figure 5a. From this analysis there are as many as 21 potential drugs for treating SCD.

**Figure 5.**
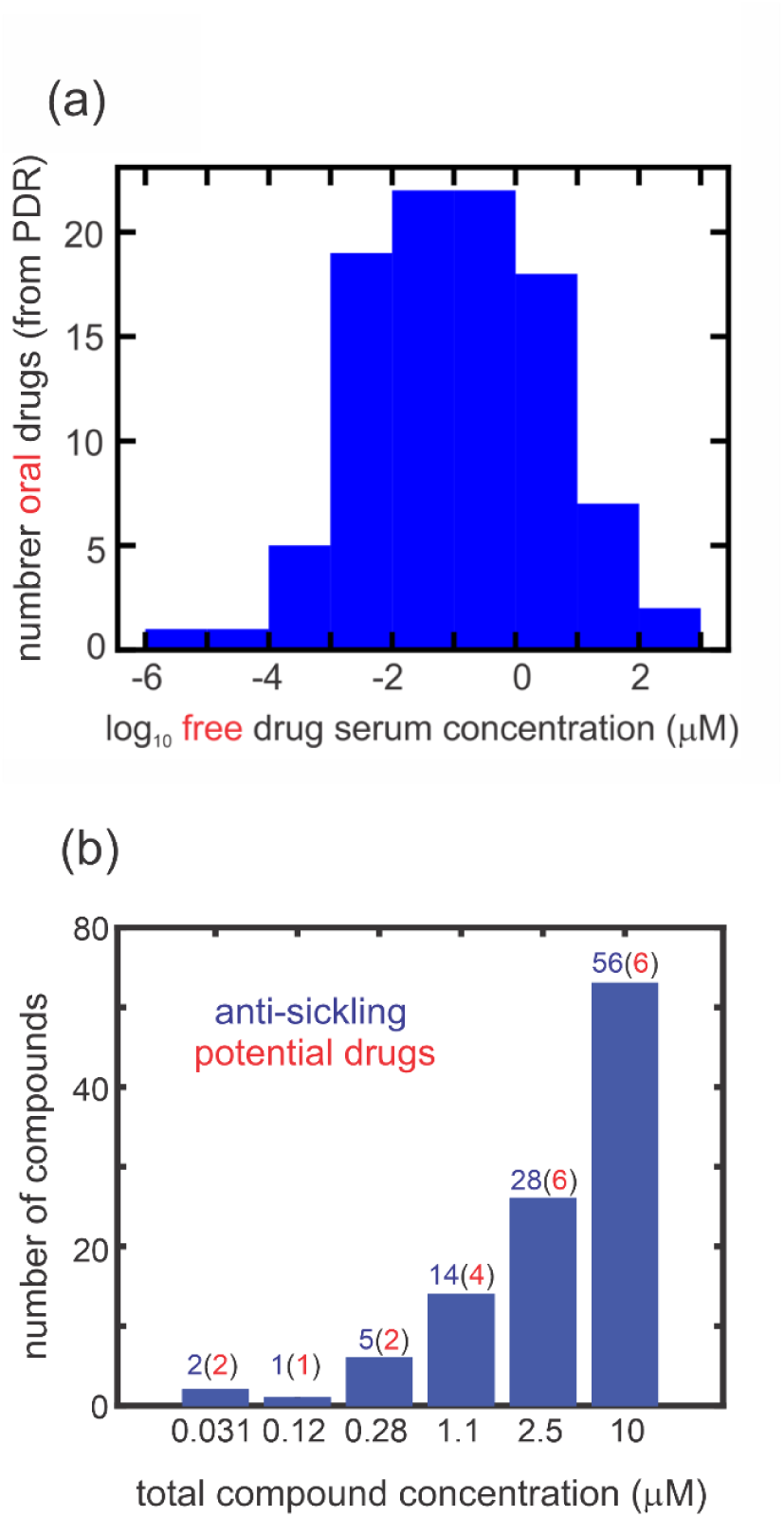
Comparing inhibitory concentrations in assay and oral drug concentrations in the PDR. (one column figure). (a) Distribution of 97 *free* oral drug concentrations (C_max_) in the 2015 version of the PDR. The free concentration was given either explicitly in the PDR or obtained from the given total serum C_max_ and the percentage bound to serum proteins. (b) Distribution of ReFrame total compound concentrations with statistically significant inhibition at each concentration, defined as 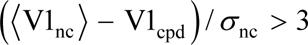. The red numbers in parentheses result from multiplying the number (blue) of compounds by the fraction of oral drugs with that free concentration or higher from the distribution in in Figure 5a. For inhibitory mechanisms other than those resulting from binding to hemoglobin, the total compound concentration is also the free concentration. For compounds that inhibit by binding to hemoglobin, the free concentration is less than the total concentration. In the 2000-fold diluted blood used for the assay, the hemoglobin molecule is at a concentration of ∼1 μM, while the molar concentration of red cells is ∼4 fM.

### Sickling curves for sickle syndromes

To assess the therapeutic potential for a given level of sickling inhibition, we compared the sickling curves for red cells from donors with various sickle syndromes. Figure 6a shows average sickling curves from deoxygenation to 5% oxygen for red cells from 65 HbSS SCD patients being treated with hydroxyurea (HU), 31 HbSS SCD patients not on HU, 9 HbSC disease patients not on HU, 9 HbSS SCD patients after HbA(T87Q) globin addition gene therapy(38, 39), an individual who is a compound heterozygote for HbS and pancellular distribution of hereditary persistence of HbF (HbS/HPFH), and from 9 individuals with sickle trait (AS). Figures 6b shows the area under the sickling curve (AUSC) for each condition (In the SVD analysis, the area under V1xS1xU1 is an accurate approximation to the AUSC). The standard deviations in Fig. 6b result from person-to-person differences, primarily due to differences in Hb composition and MCHC, with little or no contribution from experimental error. The HbS/HPFH individual and the 9 HbSS SCD patients treated with HbA(T87Q) gene addition therapy were all asymptomatic at the time of the sickling measurements.

**Figure 6.**
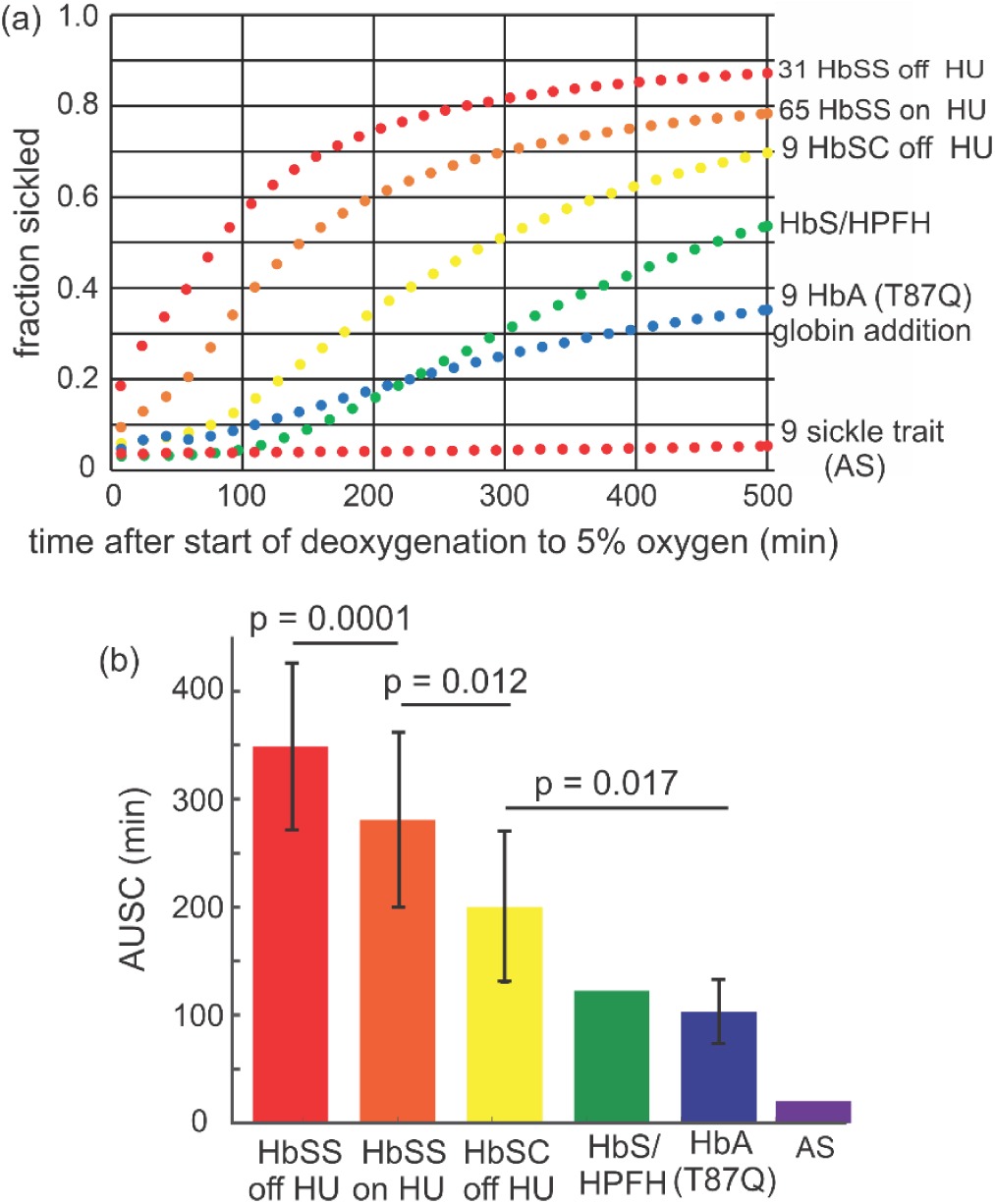
Average sickling curves for red cells from individuals with various sickle syndromes. (one column figure). (a) Sickling curves: The condition and number of samples is given next to each curve. The average compositions and MCHC’s and standard deviations from the average for each condition are given in Table 2: (b) Histogram of average AUSCs, standard deviations, and the statistical significance probability (p-values) using an unpaired, two-tailed t-test with the program Prism. The standard deviations are almost entirely due to person-to-person difference, with very little contribution from experimental error.

**Table 2.** Average compositions and MCHC’s for blood samples of various sickle syndromes in Figure 6.

| Condition | %HbS | %HbF | %HbA2 | %HbA | %HbC | %HbA(T87Q) | MCHC (g/dL) |
| --- | --- | --- | --- | --- | --- | --- | --- |
| HbSS off HU | 88.1 ± 6.7 | 7.9 ± 7.2 | 3.7 ± 0.8 | 0 | 0 | 0 | 35.0 ± 1.5 |
| HbSS on HU | 80.0 ± 8.1 | 16.3 ± 8.2 | 4.7 ± 9.9 | 0 | 0 | 0 | 35.1 ± 1.4 |
| HbSC disease | 48.4 ± 1.6 | 2.1 ± 4.0 | 4.0 ± 0.3 | 0 | 45.6 ± 2.4 | 0 | 36.1 ± 1.0 |
| HbS/HPFH | 60.2 | 38.0 | 1.8 | 0 | 0 | 0 | 35.8 |
| HbSS/HbA(T87Q) | 44.8 ± 10.2 | 2.9 ± 3.5 | 2.8 ± 0.5 | 0 | 0 | 44.5 ± 5.2 | 33.8 ± 0.61 |
| Sickle trait (AS) | 37.5 ± 3.6 | 0.3 ± 0.6 | 3.3 ± 0.2 | 59.1 ± 3.1 | 0 | 0 | 33.8 ± 0.35 |

## Discussion

The assay used in this screen must be considered quite relevant since HbS polymerization and consequent red cell sickling is the root cause of the pathology in sickle cell disease. It is also a relatively high throughput assay, as evidenced by the fact that about 80,000 sickling curves were measured in the screen, each one consisting of evaluating images of red cells at 50 times points for 100-300 cells in each well of 210 plates containing 384 wells for a total of more than 400 million red cell images. The screen was carried out in two stages. In the first stage, sickling curves were measured in triplicate for all 12,657 compounds at a concentration of 10 μM, which were incubated for 3 hrs prior to the start of deoxygenation. In the second stage, dose response measurements were made at concentrations from 1 nM to 10 μM for 3 different donors for compounds that were judged to be anti-sickling from the first stage. Since we had previously shown in our sickling study of the ionophores monensin and gramicidin that compounds can increase cell volume extremely slowly(14), we list in Table S2 of the S.I. 20 compounds that caused lysis at 10 μM but did not inhibit at a lower concentration. These compounds should be re-investigated in the future with 24 hr incubations. For the lowest inhibitory concentration of the 106 compounds listed in Tables 1 and S1, 59 inhibited sickling of cells from all three trait donors, while 47 inhibited sickling in two of three donors. As is common in research, because of failed experiments and flaws in our first analysis of the massive amount of data, the course of the experiments did not follow the original plan but took a somewhat tortuous path to finalizing the number of anti-sickling compounds. The detailed path is given in the S.I.

A critical factor for assessing therapeutic potential is to compare concentrations of anti-sickling compounds in our assay with serum concentrations in humans. To obtain an estimate of how many of the 106 anti-sickling compounds in Tables 1 and S1 are expected to be potential drugs based on their inhibitory concentration (Figure 5b), we searched the Physician’s Desk Reference (PDR) for the free concentrations (i.e., not bound to serum proteins) of oral drugs (Figure 5a). Assuming that the distribution of free serum concentrations of the 106 anti-sickling compounds discovered in our screen is the same as the distribution in the PDR, this analysis suggests that as many as 21 compounds inhibit sickling at concentrations similar to or lower than the free concentrations observed in the plasma following oral administration. These agents therefore offer potential for the treatment of sickle cell disease (Figure 5b). There are at least two caveats to using only free serum concentrations for predicting potential drugs. A drug for sickle cell disease must be taken for the lifetime of the patient and side effects that are tolerable for other patients with other diseases may not be so for sickle cell patients. Moreover, compounds may also be metabolized to a form that is less anti-sickling or even non-anti-sickling, as we discovered for the common spice curcumin, which is metabolized to a sulfate and a glucuronide (B. Metaferia *et al*., unpublished results).

For the multiple reasons described in the introduction, our screen was carried out using cells from donors with sickle trait. It was therefore important to show that compounds, which inhibit sickling of trait cells also inhibit sickling of SCD cells. Negative controls for trait cells deoxygenated to 0% oxygen and SCD cells deoxygenated to 5% oxygen to slow sickling rarely have the same amplitude and shape, so it is difficult to make a perfect comparison of inhibitory effects of a test compound for the 2 conditions. Nevertheless, comparing sickling curves for samples from SCD and trait individuals in Figure 4 strongly support the conclusion that inhibition is very similar. We might have expected larger differences than observed in Figure 4, since SCD cells are damaged from the much more frequent sickling/unsickling cycles in vivo than trait cells. The fact that they are so similar has important implications for clinical trials, for it indicates that measurements of sickling for sickle trait patients as healthy volunteers in phase 1 clinical trials for a potential anti-sickling drug should be predictive of the results for phase II trials for SCD patients.

To make a rough assessment of the expected therapeutic efficacy for a given level of sickling inhibition, we compared the sickling curves for several sickle syndromes where the final oxygen concentration was 5% (Figure 6). Given that no sickling is found for sickle trait cells with deoxygenation to 5% oxygen, the surprise was the observation of considerable sickling for cells from an asymptomatic individual with the double heterozygous condition of HbS and pancellular hereditary persistence of fetal hemoglobin (HbS/HPFH) and for cells from SCD patients effectively cured by HbA(T87Q) gene addition therapy)(36) (Q in position 87 of the γ chain of HbF is believed to be primarily responsible for reducing copolymerization of tetramers containing γ chains(40)).There is much more sickling of HbSC cells than of trait cells even though HbS is usually only about 10% higher than in trait. This has been previously explained by an increased intracellular HbS concentration in SC disease(41, 42). In our HbSC patients, the average concentration is about 5-7% higher than in trait (legend to Figure 6 and ref. (43)), which decreases the delay time 4 to 7 fold (∼1.05^30^ and ∼1.07^30^) to produce more sickling than for trait cells(7, 19, 24, 44–47). Comparing sickling curves in Figure 6 leads to the intriguing suggestion that there is a narrow range of sickling that divides asymptomatic and symptomatic individuals.

The positive news for drug development from the results in Figure 6 is that even a small reduction in sickling will be therapeutic, as observed for cells from SCD patients being treated with hydroxyurea. Moreover, the results for HbS/HPFH and HbA(T87Q) cells predict a major (even “curative”) therapeutic effect if serum concentrations that reduce sickling 2-fold (the IC50’s in Tables 1 and S1) could be achieved.

Because they lyse cells but inhibit sickling at lower concentration, we already know 17 compounds (indicated in Tables 1 and S1, foonote d) that belong to the class of anti-sickling mechanisms that increases cell volume to decrease the intracellular HbS concentration. The concept of decreasing the intracellular HbS concentration followed immediately on the decades-old observation that the delay time prior to detectable fiber formation depends on a very high power of the HbS concentration – the 30^th^ power in the initial experiments, which remains the highest concentration dependence ever observed for any molecular process. It also motivated approaches to treat the disease by decreasing the intracellular HbS concentration by means other than by increasing red cell volume(14, 48), such as introducing an iron deficiency anemia. There are multiple way of making cells iron deficient, including iron restriction, the use of chelating agents, inhibiting the enzymes that catalyze heme synthesis, and even what might be regarded as the heretical method of phlebotomy(7, 27, 49) -the method employed by physicians to treat illnesses 2000 years ago, most notably the Greek Hippocrates and Roman Galen, who subscribed to the theory that disease was caused by too much blood, one of the four humors.

To accurately determine the detailed inhibitory mechanisms for the 106 anti-sickling compounds and, more importantly, the partitioning of each of them between serum and red cells for comparing inhibitory concentrations in vitro with achievable serum concentrations in vivo will require several person-years of research.^*^ It therefore seems important to publish all of the results we have so far, which should motivate others to develop one or more of the 106 anti-sickling compounds discovered in this study into urgently needed, inexpensive drugs for treating sickle cell disease.^†,‡^ Although our primary goal is to discover an inexpensive pill, we should also consider the possibility that some of these compounds could be administered intravenously to abort or ameliorate a sickle cell crisis.

## Materials and Methods

### Sickling assay

Blood samples were collected in EDTA from anonymized donors with sickle trait according to NIH protocol 08-DK-0004 approved by the National Institute of Diabetes and Digestive Diseases Institutional Review Board. Blood samples from patients with sickle cell disease (SCD-HbSS, HbSC and HbS/HPFH) in steady-state were collected in accordance with protocol 18-H-0146 / NCT03685721 or 03-H-015 / NCT00047996 or SCD patients treated by HbA(T87Q) global addition therapy in accordance with protocols 14-H-0155 and 20-H-0141; these protocols were approved by the National Heart, Lung, and Blood Institute Institutional Review Board. The blood from sickle trait donors was diluted 2000-fold into a phosphate-buffered saline (PBS) solution, while the blood from SCD patients was diluted 1000-fold. The PBS consisted of 20 millimolar sodium/potassium phosphate, 155 millimolar sodium chloride, 300 milliosmolar, pH 7.4, 1 mg/mL dextrose, and no albumin or other protein. 10 μL of these dilutions in every well resulted in 100-300 cells in the image field for the red cells that had settled to the bottom of the wells of a polypropylene Corning 384 well plate (part no. 3770, Corning, Inc. Corning, NY). To improve mixing with the test compound, the plates were shaken on the Compact Digital Microplate Shaker (Thermo Scientific) for 1 min. The well plate was then inserted into the humidified chamber at 37°C of an Agilent Lionheart FX Automated Microscope (Agilent Technologies, Santa Clara, CA). A band pass filter (Thorlabs, FWHM = 10 ± 2 nm) at 430 nm, the peak of the deoxyhbS absorption spectrum, was added to the light path of the instrument for filtering the white light source to improve image contrast. The flow of nitrogen from the building liquid nitrogen source into the chamber containing the well plate was controlled by the Agilent gas controller (part no. 1210013). Zero percent nitrogen for trait cells or 5% oxygen for SCD cells inside the chamber was reached in 20-30 minutes. At 5% oxygen sickling of SCD cells is sufficiently slowed because of the decreased supersaturation(52) that deoxygenation is not rate limiting for sickling. Prior to preparing the well pale for SCD cells, the blood was exposed to 100% oxygen at ice temperature to melt any residual polymer that would eliminate the delay period (53). After a 3 hr incubation period to allow for the test compound to bind to its target, deoxygenation was started and continued for 10-12 hrs with images of cells collected every 15 minutes.

For the initial screen of all 12,657 compounds, wells were spotted by Calibr with 100 picomoles of each compound to give a concentration of 10 μM after the addition of 10 μL of the dilute red cell suspension. For dose response measurements, the wells contained from 10 femtomoles to 100 picomoles spaced at approximately factors of three intervals to give compound concentrations from ∼1 nM to 10 μM. Triplicate plates for both the initial screen and dose-response measurements were provided by Calibr in order to assess differences in the inhibitory response for trait cells from three different donors. Cells containing no test compound in wells of two columns of the plate served as negative controls; cells in wells of two columns in 170 milli-osmolar PBS and containing no test compound served as positive controls. The hypo-osmotic effect of 170 milli-osmolar swells cells and often so much so that the cells are sphered, which increases the cell volume by a factor of ∼1.4.(54) Swelling reduces the intracellular hemoglobin concentration sufficiently to completely prevent sickling for 12 hrs or more because of the enormous sensitivity of the polymerization kinetics to the hemoglobin concentration (the 30^th^ power concentration dependence results in a 30,000-fold increase in the delay time for a 1.4-fold decrease in concentration(19)).

In comparing sickling for sickle trait and SCD cells in Figure 4, all 14 compounds were purchased. The sources were: curcumin (Sigma), voxelotor (MedChem), mavatrep (MedChem), entrectinib (MedChem), pimobedan (MedChem), calcimycin (Medchem), laflunimus (MedChem), MI-773 (MedChem), COH-29 (Medchem), fursultiamine (MedChem), WP-1066 (MedChem), NSC-150117 (MedChem), etacrynic acid (Sigma), phanquinone (Sigma), mitapivat (SelleckChem).

### Image analysis

The image analysis is based on the fact that fiber formation is the only possible cause of a change in red cell morphology at optical microscope resolution, apart from rare fluctuations that tilt red cells onto an edge that gives an elongated projection counted as sickled; it is strongly supported by the observation in single SCD cells that changes in cell morphology and scattered light caused by polymerization are simultaneous (53) and the images of cell morphology shown to contain polymerized HbS from measurements of oxygen binding for individual cells(55, 56). The most important metrics for detecting a morphologic change are the loss of roundness, the loss of a more transparent center, and the decrease in the projected area of the cell. Two kinds of analysis were performed. In one, the sickling time was determined as the time when at least two of the three metrics of the cell image changed in the appropriate direction compared to the preceding image ∼15 minutes earlier. In the second, machine learning was employed, in which a model was constructed from the assignment by an experienced observer of about 1,000 cells as either sickled or not sickled based on the above 3 metrics (see ref. 5 for more details). Machine learning is particularly useful for analyzing images of SCD cells, since cells in the very first image when there is no deoxygenation, so-called irreversibly sickle cells (ISC’s), are counted as sickled, while in the first analysis they would not be counted because the shape of ISC’s may not change at all when fibers form during the 10-12 hr course of the experiment. The output of an experiment is a plot of the fraction sickled versus time for cells incubated with each library compound and the average fraction sickled versus time and standard deviation from the average fraction at each time point for the negative and positive controls.

### Singular value decomposition of sickling curves

For analysis of the sickling curves, the powerful matrix-analytic method of singular value decomposition (SVD) of the data was employed(37). The data consist of the fraction sickled at 50 times points following the start of deoxygenation. The data matrix, **D** is a 50×384 matrix. SVD exactly reproduces **D**, including the noise, with a product of three matrices, **UxSxV^T^**. **U** is a 50×50 orthogonal matrix of the fraction sickled versus time at 50 time points for the images in each of the 384 wells, **S** is a 50×384 matrix with nonzero elements (the so-called singular values) only on the principal diagonal that is a measure of the contribution of each component of **U** to **D**, and **V** is a 50×384 matrix the columns of which are the amplitudes for each component of **U**. All but the first 2 or 3 components of the SVD represent primarily noise. As judged by the magnitude of the singular values, the complete set of 384 sickling curves on a single well plate could therefore be sufficiently well represented noise-free by just the first component of the SVD, i.e. S1xU1xV1. If, for a particular concentration of the test compound, 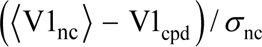, the average V1 for the negative control wells minus V1 for the test compound well, divided by *σ*_nc_, the standard deviation from the average V1 for the control wells, is positive and larger than 3, it was considered anti-sickling. The standard deviation of V1 is a single number and is therefore independent of time, explaining the difference in the time dependence between the standard deviations in the raw sickling curves and the sickling curves resulting from the SVD. Because sickling curves for the negative controls vary with row number, presumably because of small differences in the rate of deoxygenation hemoglobin inside the red cells sitting at the bottom of the well, the values of V1 for the row of the test compound plus the V1’s for negative controls from 2 adjacent rows were used to obtain 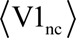, apart from rows 2 and 15, in which case the V1’s from only 1 adjacent row was used to calculate 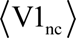. To improve the accuracy of 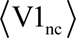, the V1’s for sickling curves from an additional 12 (or 8) wells that contain 1, 3 and 10 nM of the test compound were also used as negative controls, since no anti-sickling was observed for any of the compounds at these concentrations (Table 1).

Even though the concentration of HbS in the hypo-osmolar buffer of the positive controls is too low for any polymerization to occur, the image analysis algorithm invariably counted a small fraction as sickled because of changes over the hours duration of the assay. Since such changes could also occur for cells incubated with test compounds, the percent inhibition of sickling in the wells containing test compounds was defined as:

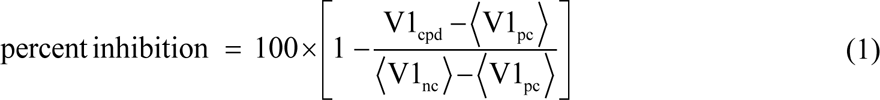

### Fitting dose response data with smooth curve

The model used to provide a least squares description of the dose response data is very similar to the log-logistic model proposed in equation 2 of the paper by Ritz et al.(57)

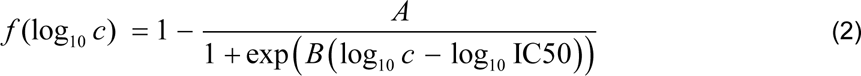

where *c* is the compound concentration. The 3 adjustable parameters are *A*, *B*, and log_10_ IC50, the concentration in micromolar at 50% inhibition. An assumption in using this equation is that inhibition is complete, *f* (log *c*) = 1 at infinite compound concentration. Constraints were applied to keep the value of the function close to 0 at the lowest compound concentrations and to insure that the fitted curves have a sigmoid shape. The sum of squares (*ssq*) was minimized with the constraint terms *f*1 and *f*2 determined empirically.

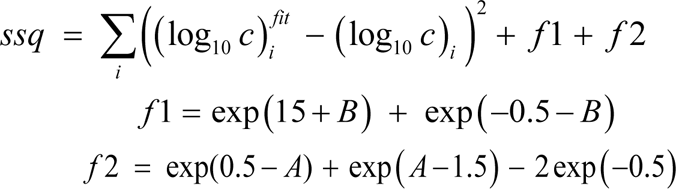

The term *f*1 constrains the curve to have a sigmoid shape with the adjustable parameter, *B*. The term *f*2 is an additional constraint with the adjustable parameter of *A*. In the log-logistic model, *A* is the difference between the upper and lower asymptotes of the response, which in the SVD analysis is the difference between 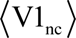 and 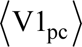. At concentrations too low to be inhibitory, (S1× U1× V1)_cpd_ can be larger than 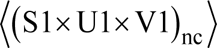, which produces negative deviations from 0. After fitting, differences between the fitted and experimental points (residuals) were fit to a Gaussian distribution. If a residual was greater than three standard deviations from the mean, the relevant experimental point was removed as an outlier. The data were then refit with the outliers removed.

## Acknowledgements

This work was supported by the intramural research program of the National Institute of Diabetes and Digestive and Kidney Diseases (NIDDK) and the National Heart, Lung, and Blood Institute of the National Institutes of Health (NHLBI). We thank H. Franklin Bunn (Harvard Medical School) for advice throughout this project and for a careful reading and comments on the manuscript, Rafael Daniel Camerini-Otero (NIDDK) for suggesting that W.A.E. work on drug development, Marc Reitman (NIDDK) for encouraging us to write this article at this stage of our drug search, and Diane Kambach of Agilent technologies for her technical help in adapting the Lionheart for the transmission measurements with nitrogen deoxygenation. We also thank the Scripps California Institute for Biomedical Research (Calibr) for their gift of the entire ReFrame library and Kaycie Morwood and Emily Chen at Calibr for preparing the well plates containing the library compounds. We thank David Wood and Jonathan Sachs (University of Minnesota) for a preprint of their article on high-throughput screening.

## Supplementary Information for

### Description of video of cells sickling

Images were collected every 15 min for 16 hrs, so the 26 second video has been sped up by about a factor of ∼2,000. The contrast increase early in the video is due to the change in the absorption spectrum of hemoglobin from oxy (peak at 415 nm) to deoxy (peak at 430 nm, the center of the band pass filter).

**Table S1:**
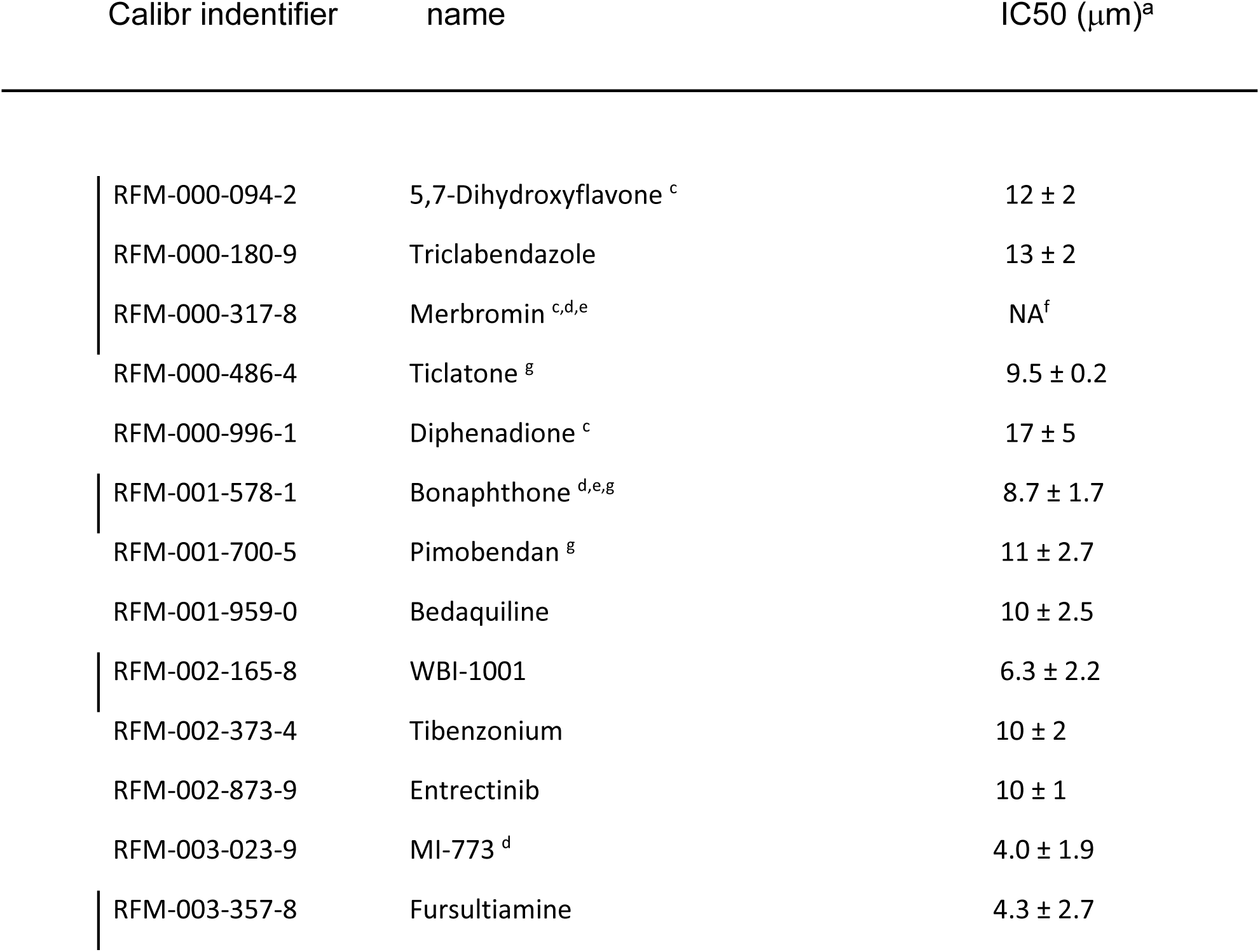
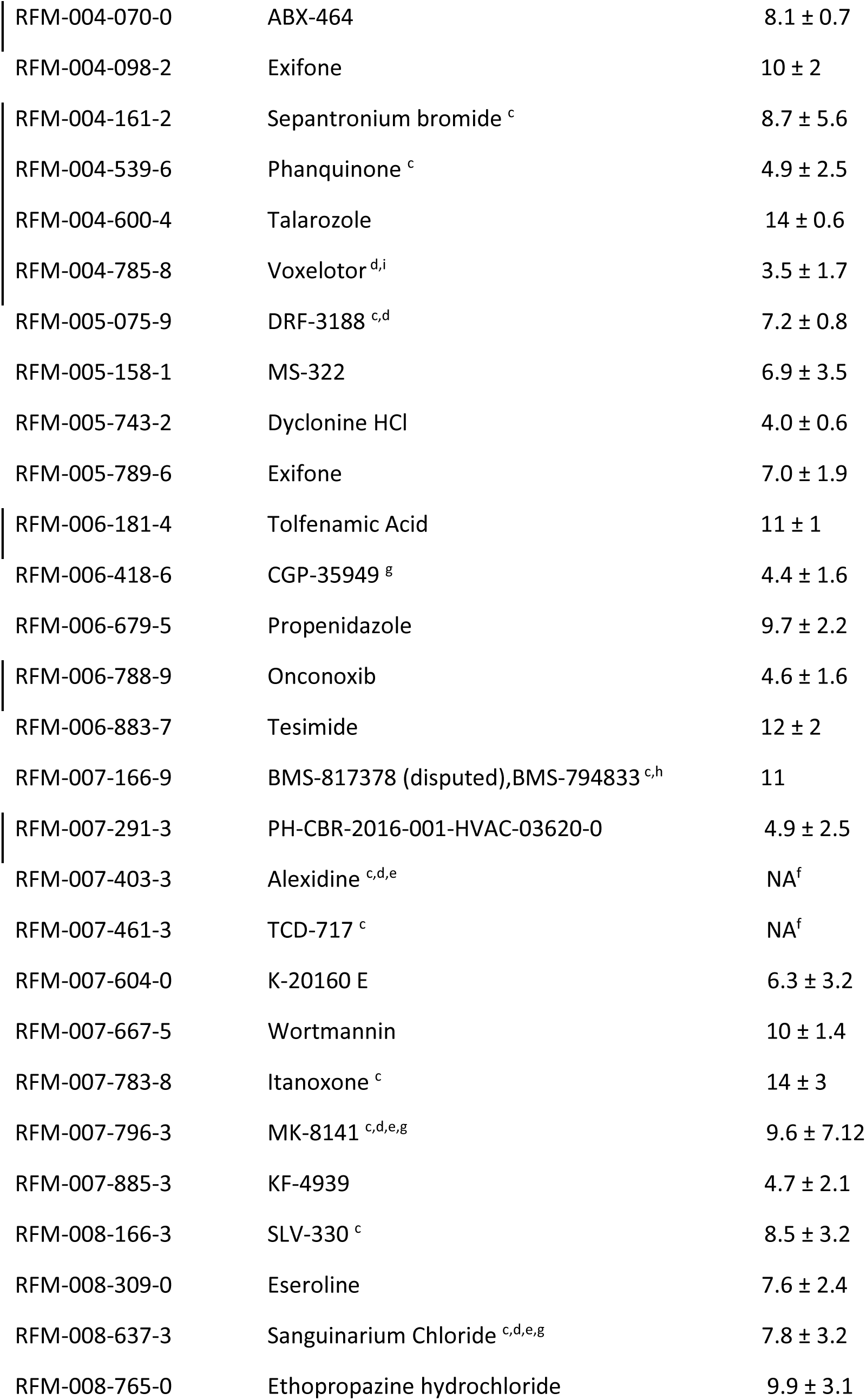
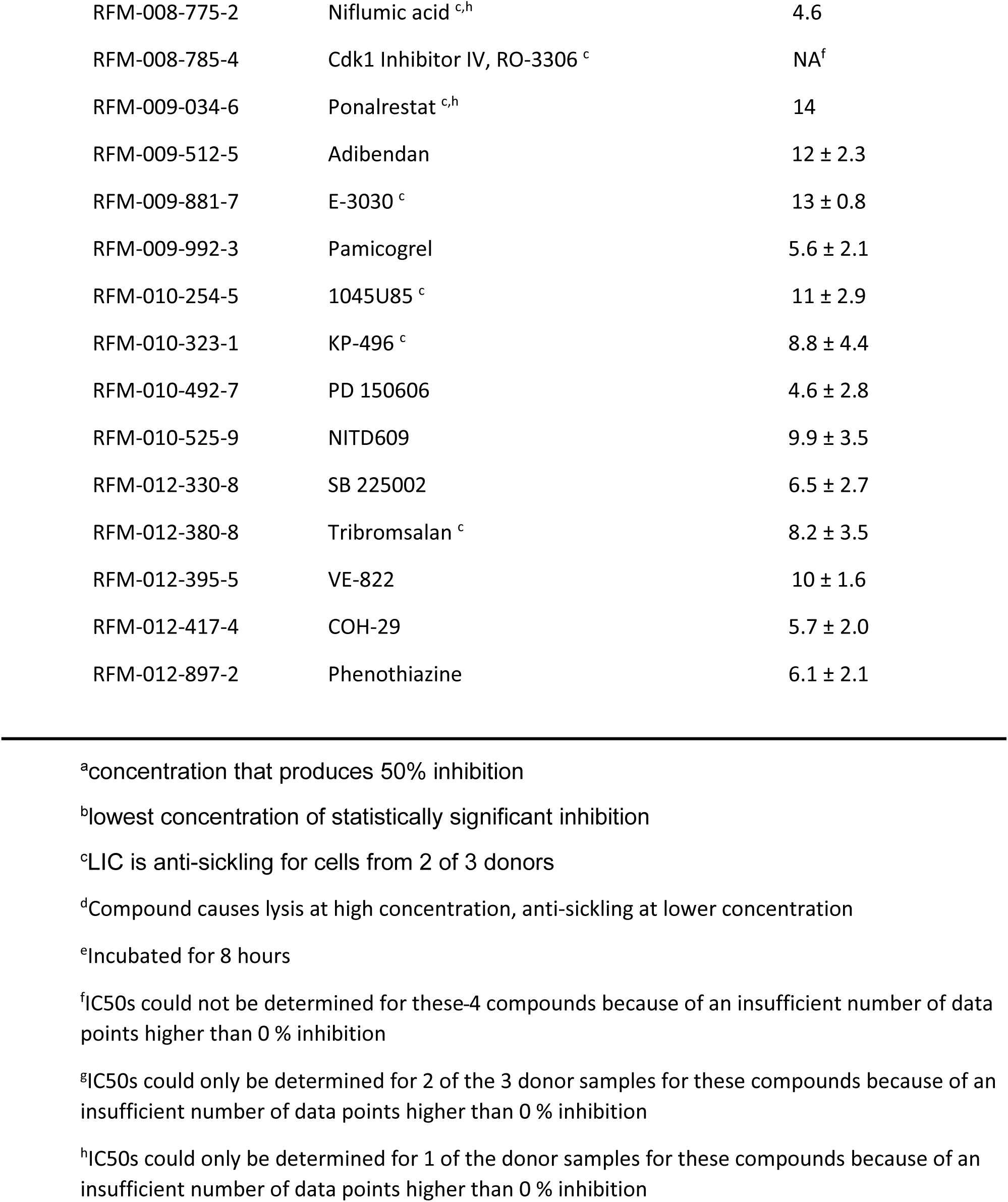
List of compounds that inhibit sickling at 10 μm^b^ and do not inhibit at lower concentrations.

## Additional details of screen

### Organization of well plates

Corning 384 well plates were shipped to Calibr at Scripps Institute in La Jolla, CA and returned to NIH after spotting 100 picomoles of all 12,657 compounds of the ReFrame library. 10 μL of the sickle trait red cell suspension was added to give a compound concentration of 10 μM. Compounds that reduced sickling were then studied in a second set of experiments that measured the percent inhibition as a function of the compound concentration (i.e., dose response measurements). For these experiments, 9 wells were spotted with 9 concentrations of a compound from 10 femtomoles to 100 picomoles spaced at approximately factors of 3 intervals to produce final concentrations from ∼1 nM to 10 μM. Triplicate plates for both the initial screen and dose-response measurements were sent by Calibr in order to assess variation in the inhibitory response from 3 different donors. Wells in rows 2-15 and columns 4-21 of a 384 (16 rows, 24 columns) well plate were spotted with compounds. Cells in rows 2-15 and columns 3 and 22 containing no test compound served as negative controls. Cells in a 170 milli-osmolar PBS buffer in rows 2-15 and columns 2 and 23 containing no test compound served as positive controls. Because of imperfect homogeneity in the control of the gas atmosphere above the well plate in the humidified chamber, the rate of deoxygenation is not exactly the same for every well of the plate, with the rate decreasing with increasing row number. Consequently, only the cell suspensions for the negative and positive controls in the same row and the 2 rows adjacent to the row with the test compound were considered (or 1 adjacent row for rows 2 and 15). Also, image analysis of the cells in rows 1 and 16 and edge columns 1 and 24 were not considered because of edge effects.

### Selection of compounds for dose-response measurements

In the analysis of the first step of the screen at 10 μM, the average of the sickling curves for all 28 negative control wells on the plate were used to determine whether a compound is anti-sickling. Singular value decomposition (SVD) was used to make this determination. Using the 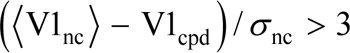 criterion from the SVD analysis for statistically significant inhibition, 335 compounds were further studied in dose-response experiments for each one at 9 compound concentrations between ∼1 nM and 10 μM, spaced by approximately factors of 3. Of the 335, 62 compounds lysed cells at 10 μM, which were pursued because swelling red cells is a viable therapeutic strategy, so it was important to study these compounds at concentrations less than the 10 μM lysis concentration. A later analysis using only control wells in the same row as the drug wells and the two adjacent rows, dose response measurements for these 335 compounds showed that 192 were false positives. The reason for this is that the sickling curve is sensitive to the rate of deoxygenation, which was later discovered to decrease in the Lionheart humidified chamber as the row number increases, presumably because of imperfect homogeneity in the percent oxygen as a function of time in the atmosphere above the well plate. Adding a cover to the well plate with an 11 by 7 array of 1.7 mm holes spaced 9.0 mm apart improved the homogeneity but did not make it perfectly homogenous. The sickling curves for 12,657 compounds were therefore completely reanalyzed with the more limited number of wells as negative controls, which also uncovered 92 false negatives, i.e., compounds with 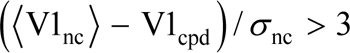.

The Lionheart microscope was not designed to be optimal for our assay. In addition to the lack of homogeneity in the time dependence of the % oxygen in the humidified chamber of the Lionheart instrument, a troublesome defect was the laser auto-focusing cube, which was not stable in many experiments. We suspect that a lens mount in the cube moves when the nitrogen flow is high, as it is at the start of the experiment, but then returns to its original position when the nitrogen flow is slowed considerably upon obtaining the final desired percent oxygen. Consequently, the first 5-10 images from each well can become badly out of focus before the focus returns to its original setting at the start of the nitrogen flow. For this reason, the dose response measurements had to be repeated on 70 of the compounds with de-focused images.

### 4 additional representative data sets

**Figure S1.**
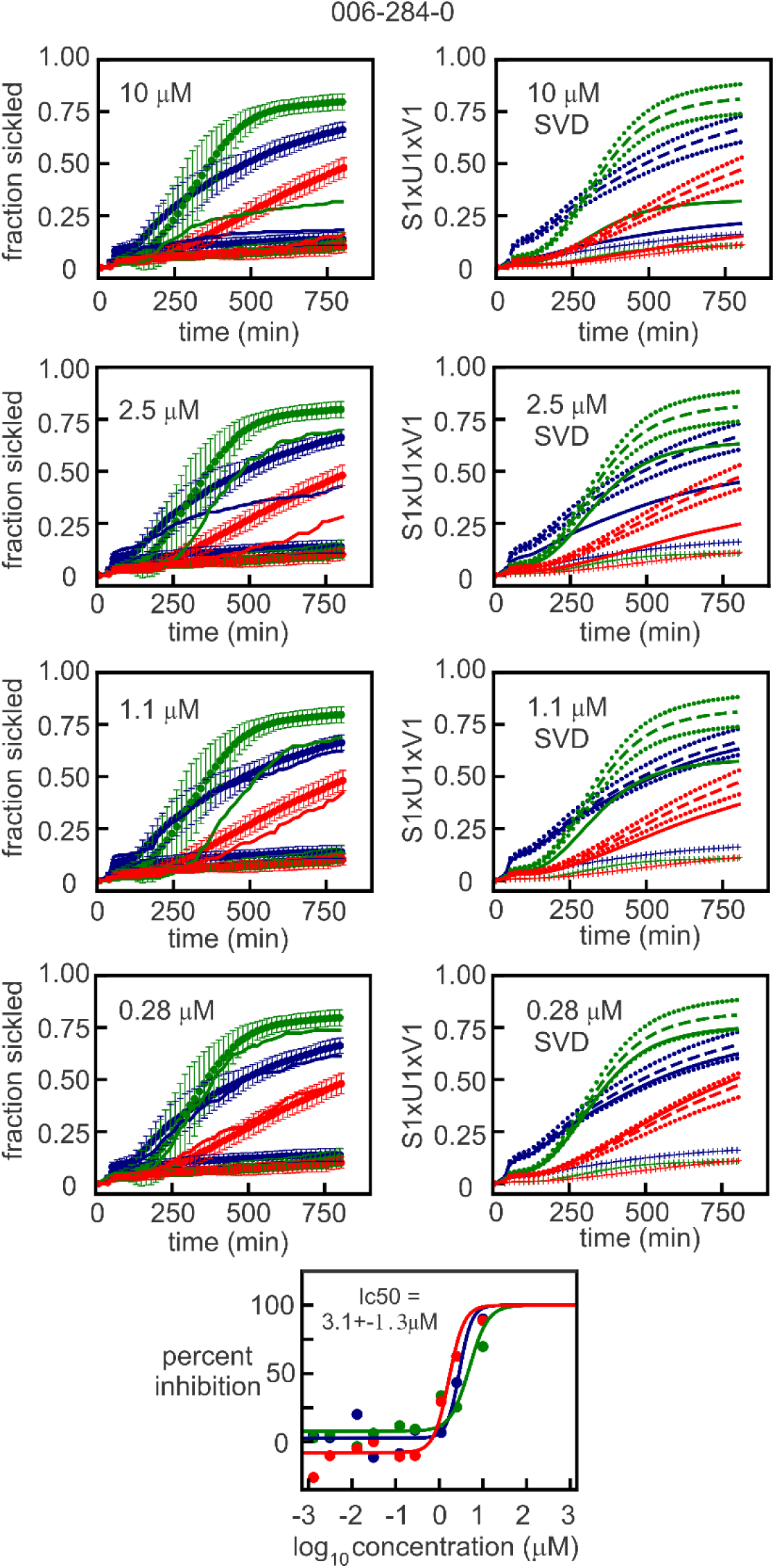
Representative dose response data for ReFrame compound TS-80. (a) Sickling curves at four different concentrations of TS-80. Panels on the left are the measured curves, while panels on the right show the curves represented by the first component of the SVD. The continuous curves in both left and right panels are the sickling curves for the single wells containing the test compound. The three colors represent samples from three different sickle trait donors with hemoglobin compositions: blue donor: 59.8 %HbA, 35.5 %HbS, 3.8 %HbA2, <1.0 %HbF, MCHC 30.0 g/dL; green donor: 59.8 %HbA, 36.4 %HbS, 3.6 %HbA2, <1.0 %HbF, MCHC 28.5 g/dL; red donor: 58.4 %HbA, 38.0 %HbS, 3.3 %HbA2, <1.0 %HbF, MCHC 26.9 g/dL. In the left panels, the upper points are the average and standard deviations from the average for six negative controls, while the lower points are the average and standard deviations from the average of six positive controls. In the right panels, the long-dashed curves are the average sickling curves (S1xU1v1)) of the negative controls with standard deviations indicated by the short-dashed curves. The lower points and vertical bars are the average and standard deviations for the positive controls. (b) Dose response data from SVD analysis. The percent inhibition is given by equation (1) in **Methods** section in the main text.

**Figure S2.**
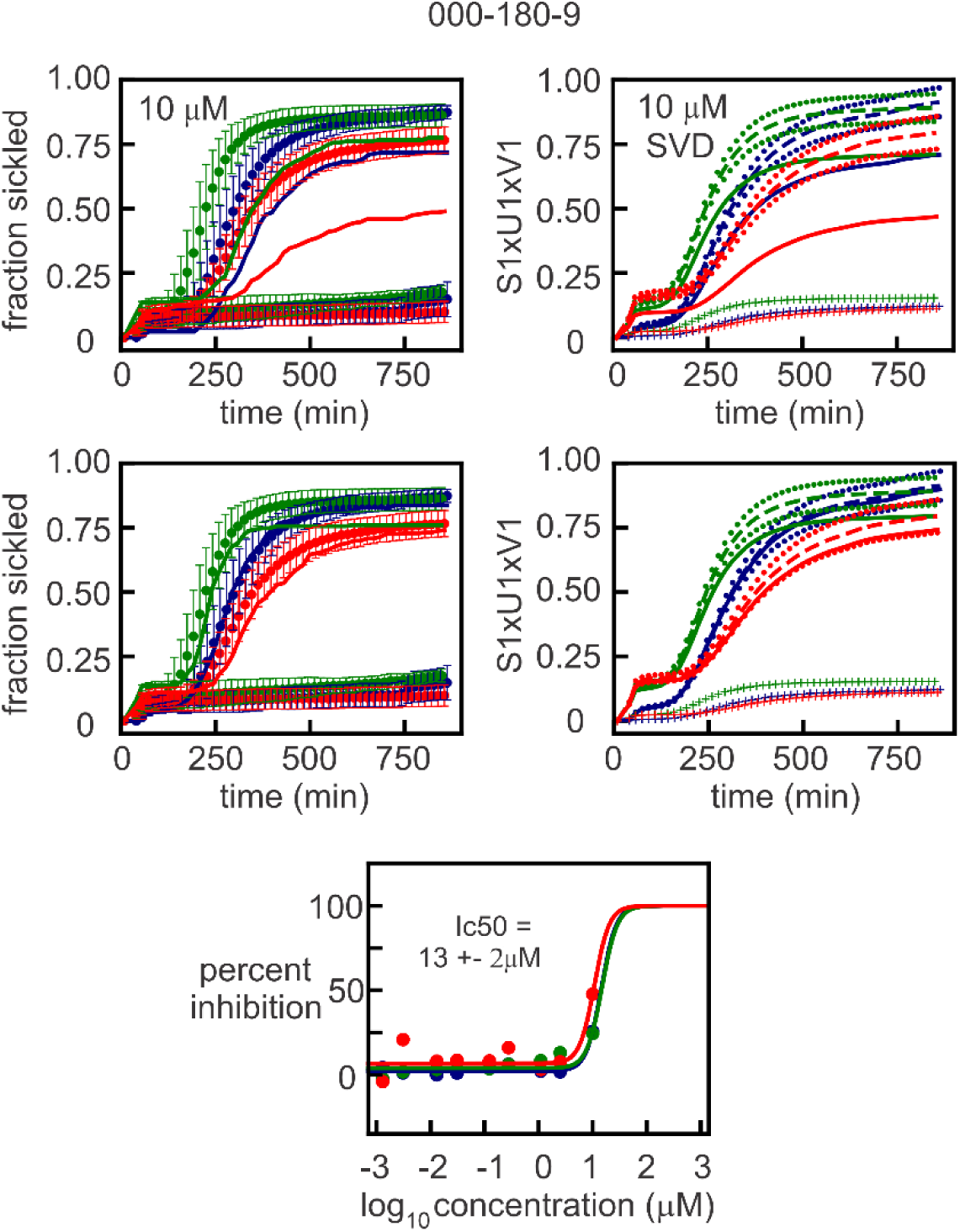
Representative dose response data for ReFrame compound triclabendaole. The hemoglobin compositions are: blue donor: 56.1 %HbA, 40 %HbS, 3.7 %HbA2, <1.0 %HbF, MCHC 29.5 g/dL; green donor: 62.7 %HbA, 33.4 %HbS, 3.5 %HbA2, <1.0 %HbF, MCHC 27.6 g/dL; red donor: 59.8 %HbA, 36.4 %HbS, 3.6 %HbA2, <1.0 %HbF, MCHC 28.5 g/dL. Remainder of legend is same as for Figure S1.

**Figure S3.**
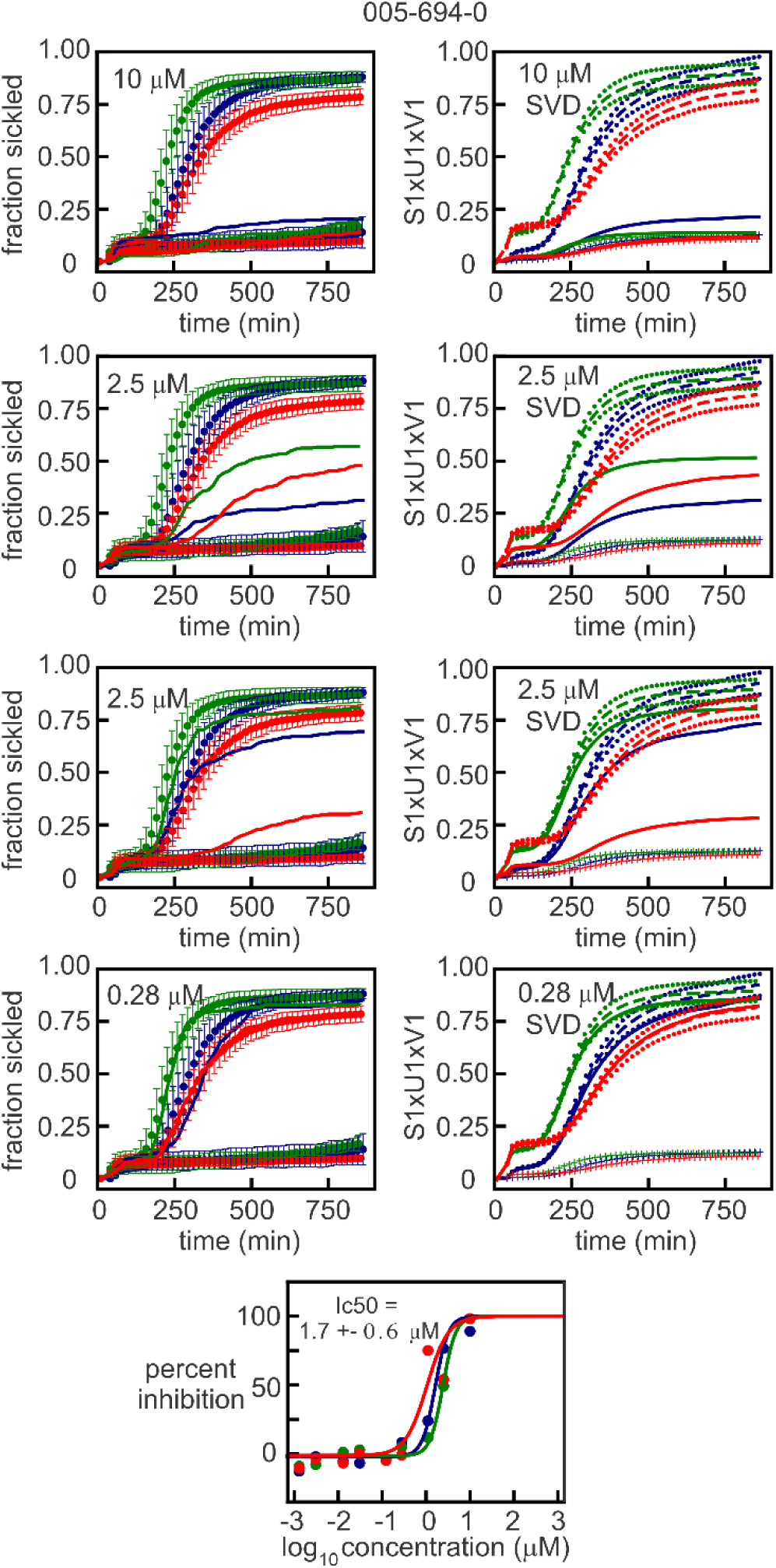
Representative dose response data for ReFrame compound cyclovalone. The hemoglobin compositions are: blue donor: 59.8 %HbA, 35.5 %HbS, 3.8 %HbA2, <1.0 %HbF, MCHC 30.0 g/dL; green donor: 59.8 %HbA, 36.4 %HbS, 3.6 %HbA2, <1.0 %HbF, MCHC 28.5 g/dL; red donor: 58.4 %HbA, 38.0 %HbS, 3.3 %HbA2, <1.0 %HbF, MCHC 26.9 g/dL. Remainder of legend is same as for Figure S1.

**Figure S4.**
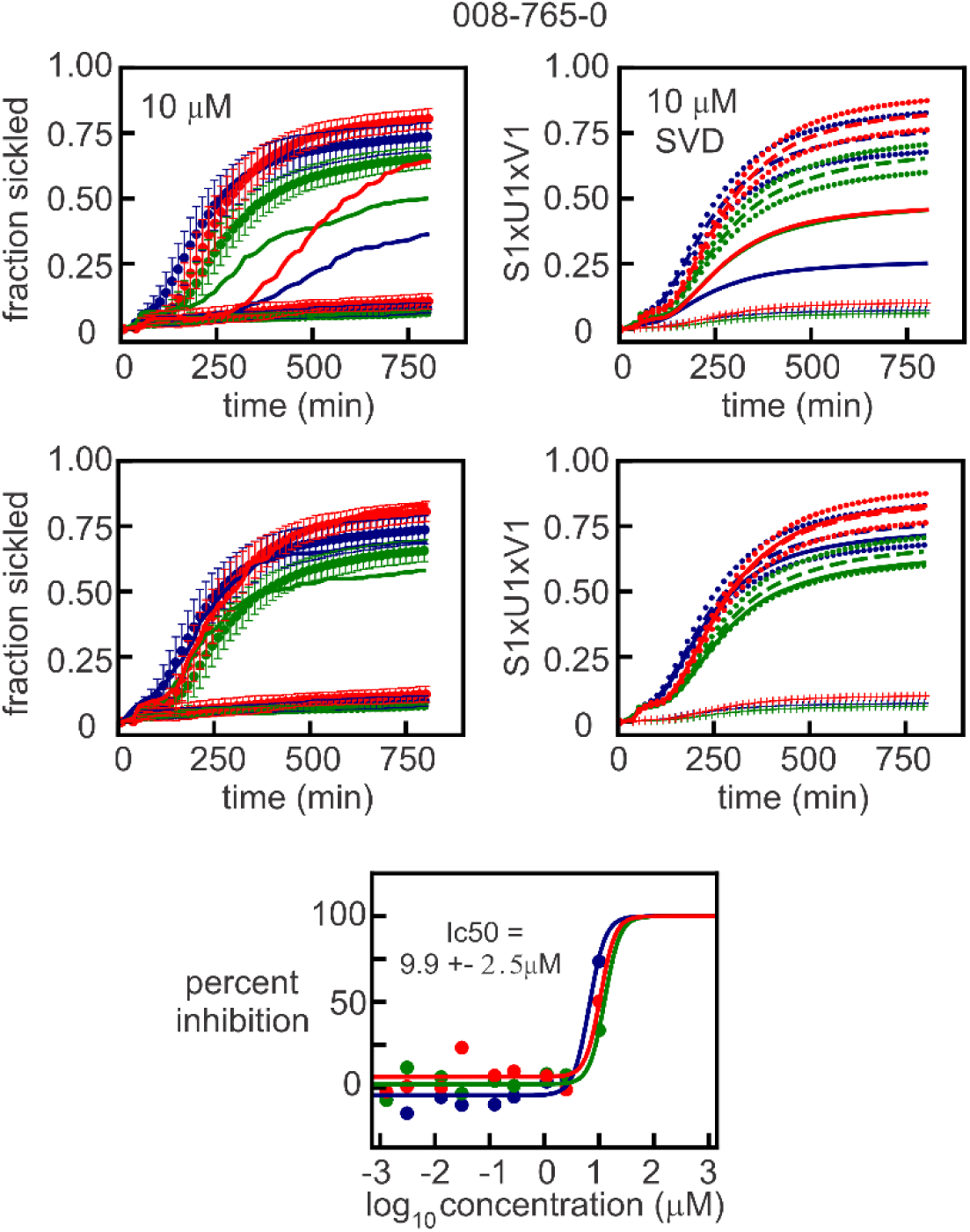
Representative dose response data for ReFrame compound ethopropazine hydrochloride. The hemoglobin compositions are: blue donor: 59.8 %HbA, 35.5 %HbS, 3.8 %HbA2, <1.0 %HbF, MCHC 30.0 g/dL; green donor: 59.7 %HbA, 37.8 %HbS, 3.8 %HbA2, <1.0 %HbF, MCHC 26.9 g/dL; red donor: 58.1 %HbA, 37.6 %HbS, 3.6 %HbA2, <1.0 %HbF, MCHC 27.8 g/dL. Remainder of legend is same as for Figure S1.

The compounds listed in Table S2 showed lysis at 10 μM but no anti-sickling at a lower concentration. They should be further investigated after 24 hrs of pre-incubation before the start of deoxygenation.

**Table S2.** **List of compounds that lyse cells at 10 μM but do not inhibit sickling at lower concentrations**

|  |  |
| --- | --- |
| RFM-000-770-5 | Alectinib Hydrochloride |
| RFM-000-900-7 | Anidulafungin |
| RFM-003-232-6 | Tannic Acid |
| RFM-003-936-1 | ST-1859 |
| RFM-004-540-9 | Ebselen |
| RFM-006-074-2 | Provecta <sup>a</sup> |
| RFM-006-443-7 | AGN-194310 |
| RFM-006-501-0 | OT 4003 |
| RFM-006-502-1 | OT 4003 <sup>a</sup> |
| RFM-006-841-7 | Mercufenol Chloride <sup>a</sup> |
| RFM-006-939-6 | R-213129 |
| RFM-007-934-5 | AGN 203818 |
| RFM-008-420-8 | Xanthohumol |
| RFM-010-146-2 | Cethexonium Bromide |
| RFM-010-391-3 | Mecetronium Ethylsulfate |
| RFM-010-455-2 | Fendosal |
| RFM-011-755-5 | PLD-147 |
| RFM-012-549-5 | Purvalanol A |
| RFM-012-595-1 | Mercuric Chloride |
| RFM-012-876-7 | Elacestrant (dihydrochloride) |

<sup>a</sup>Incubated for 8 hours

The following pages are the dose response plots for 99 of the 106 anti-sickling compounds

(see footnote dTable 1 of main text)

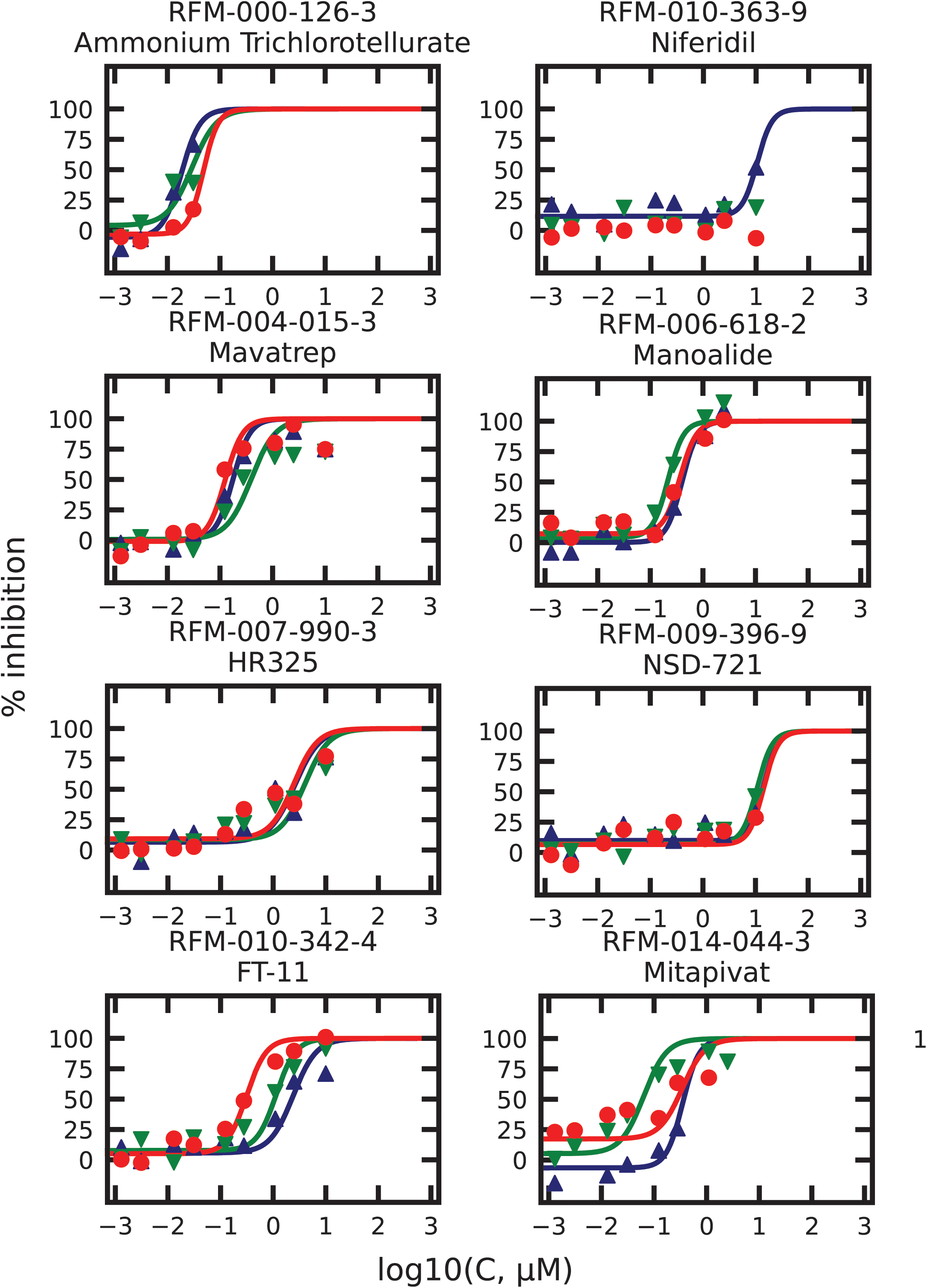
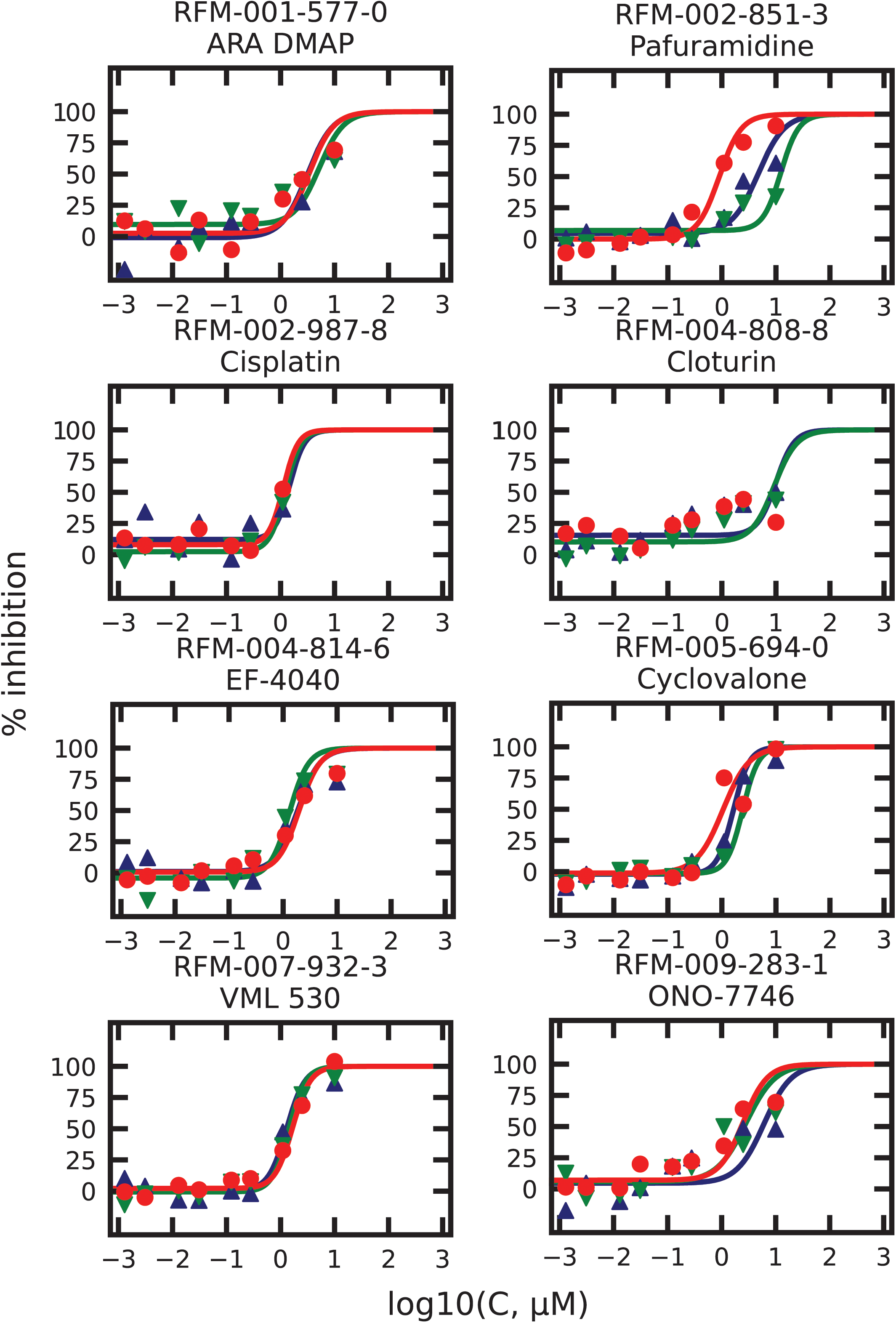
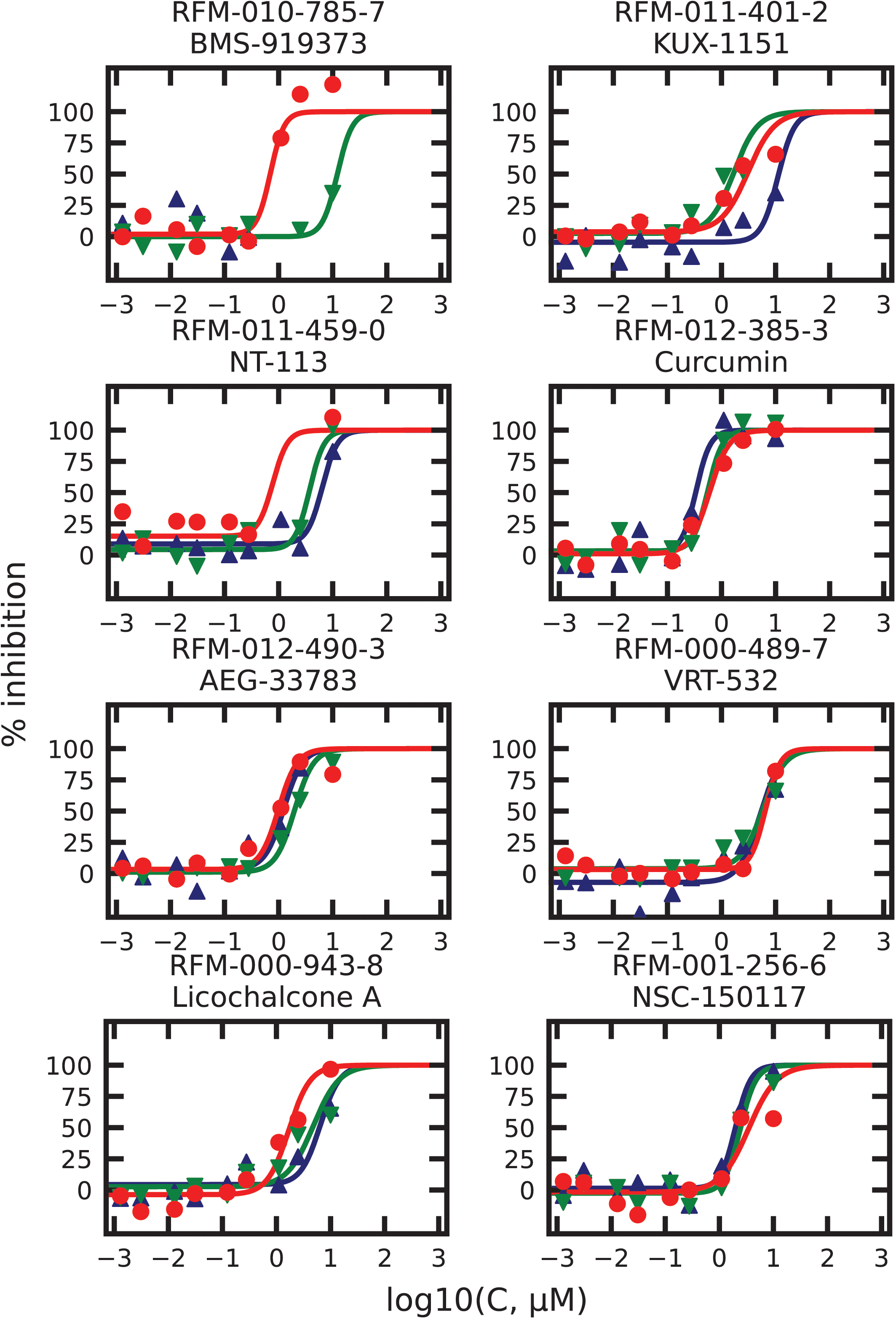
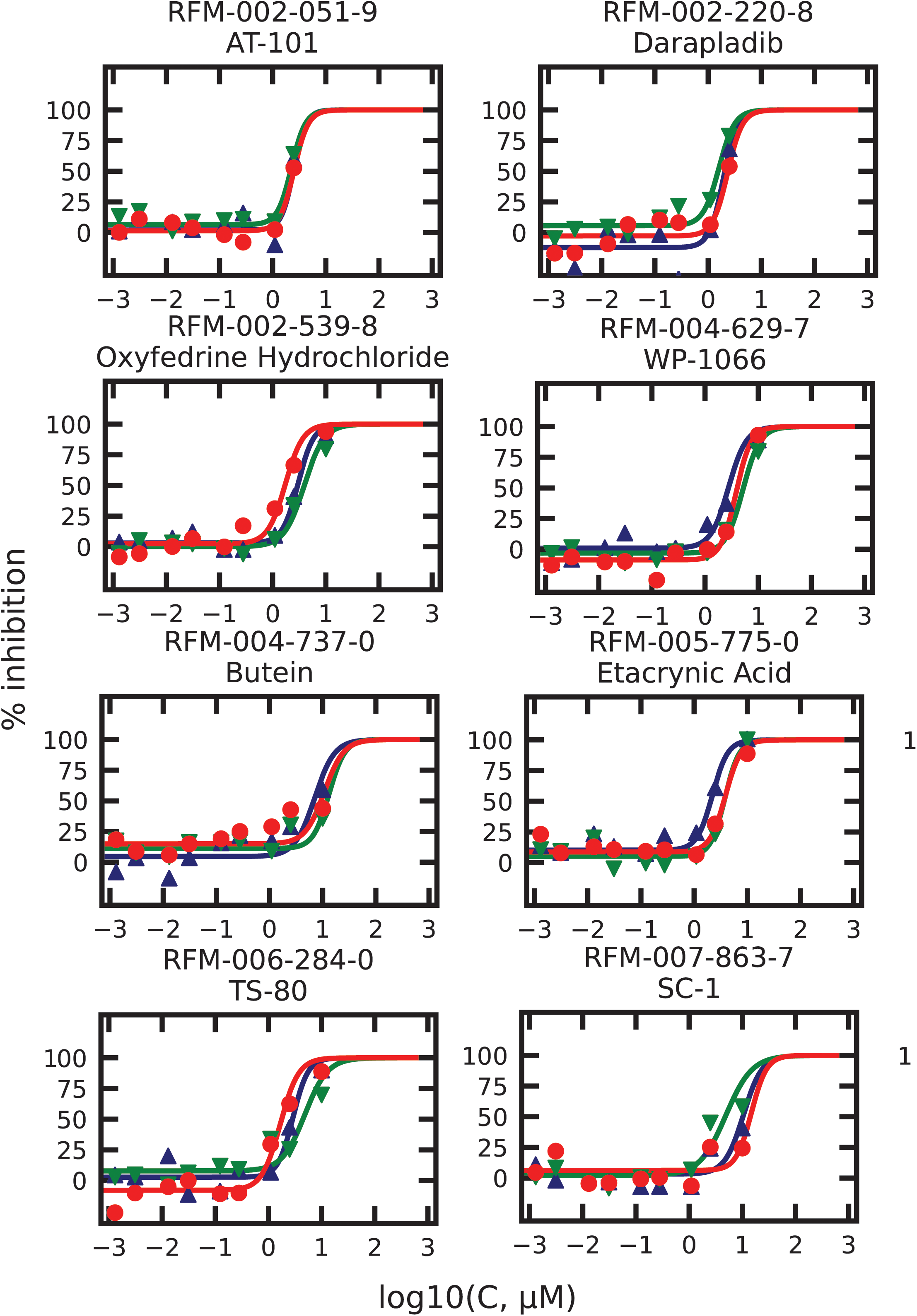
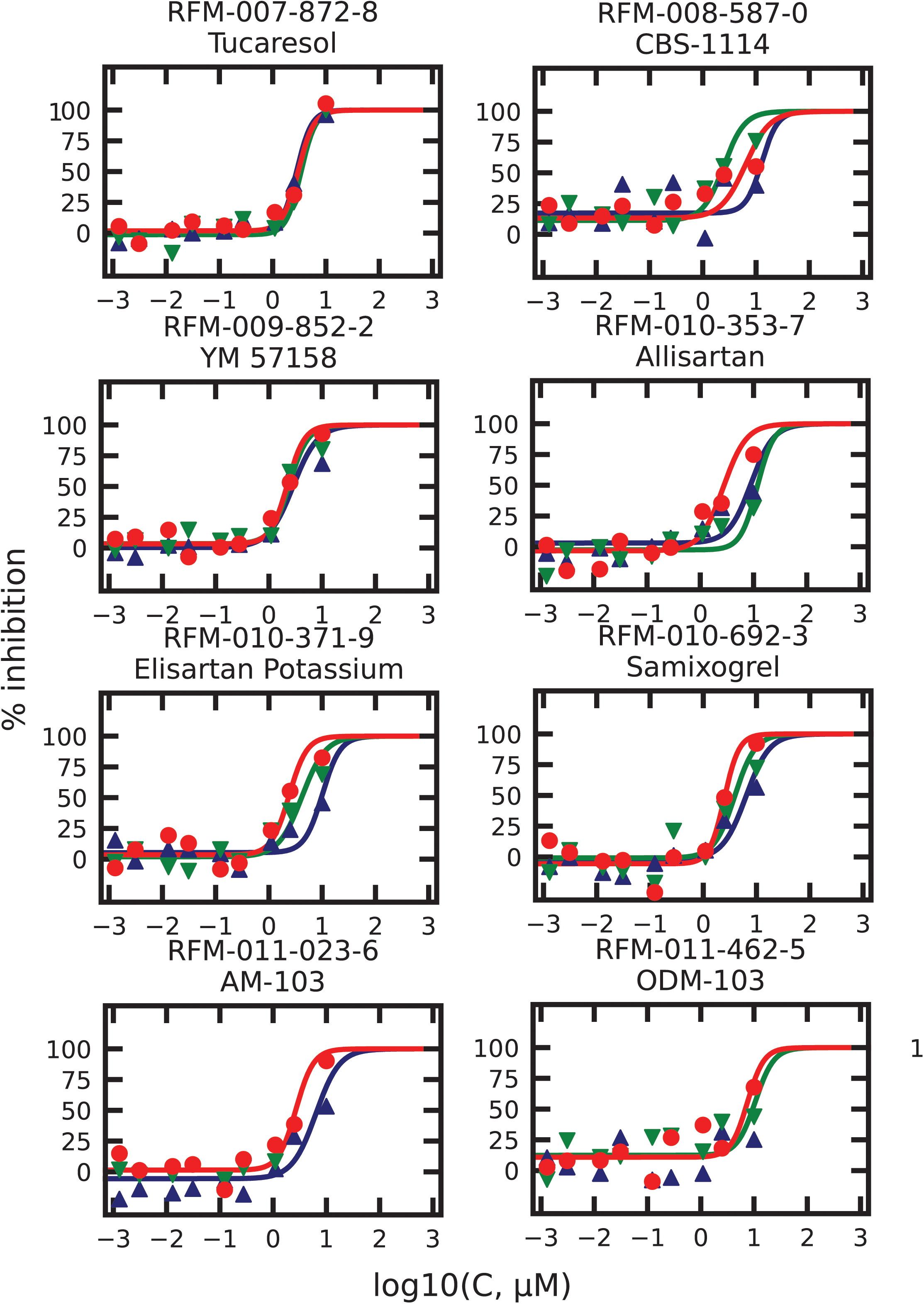
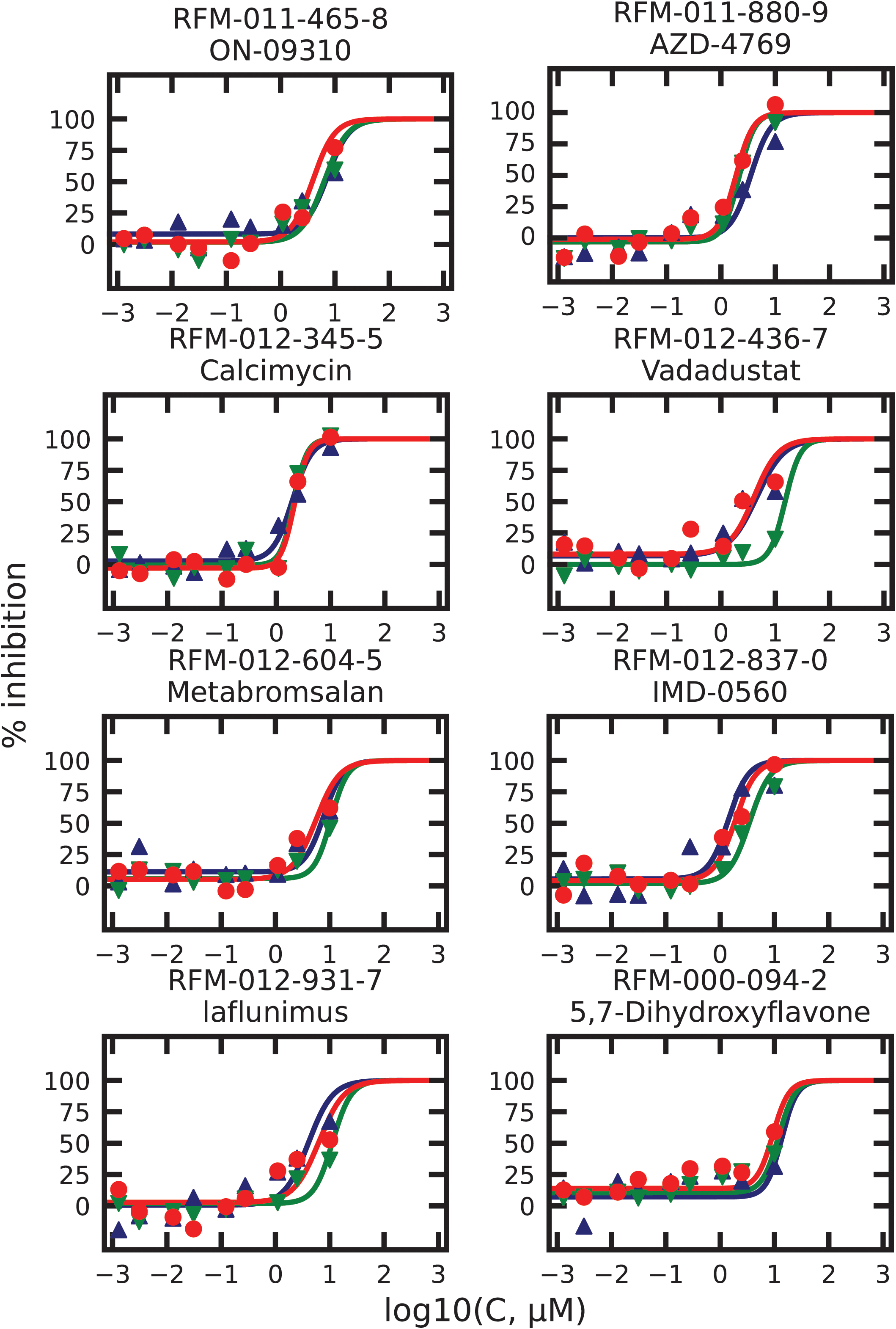
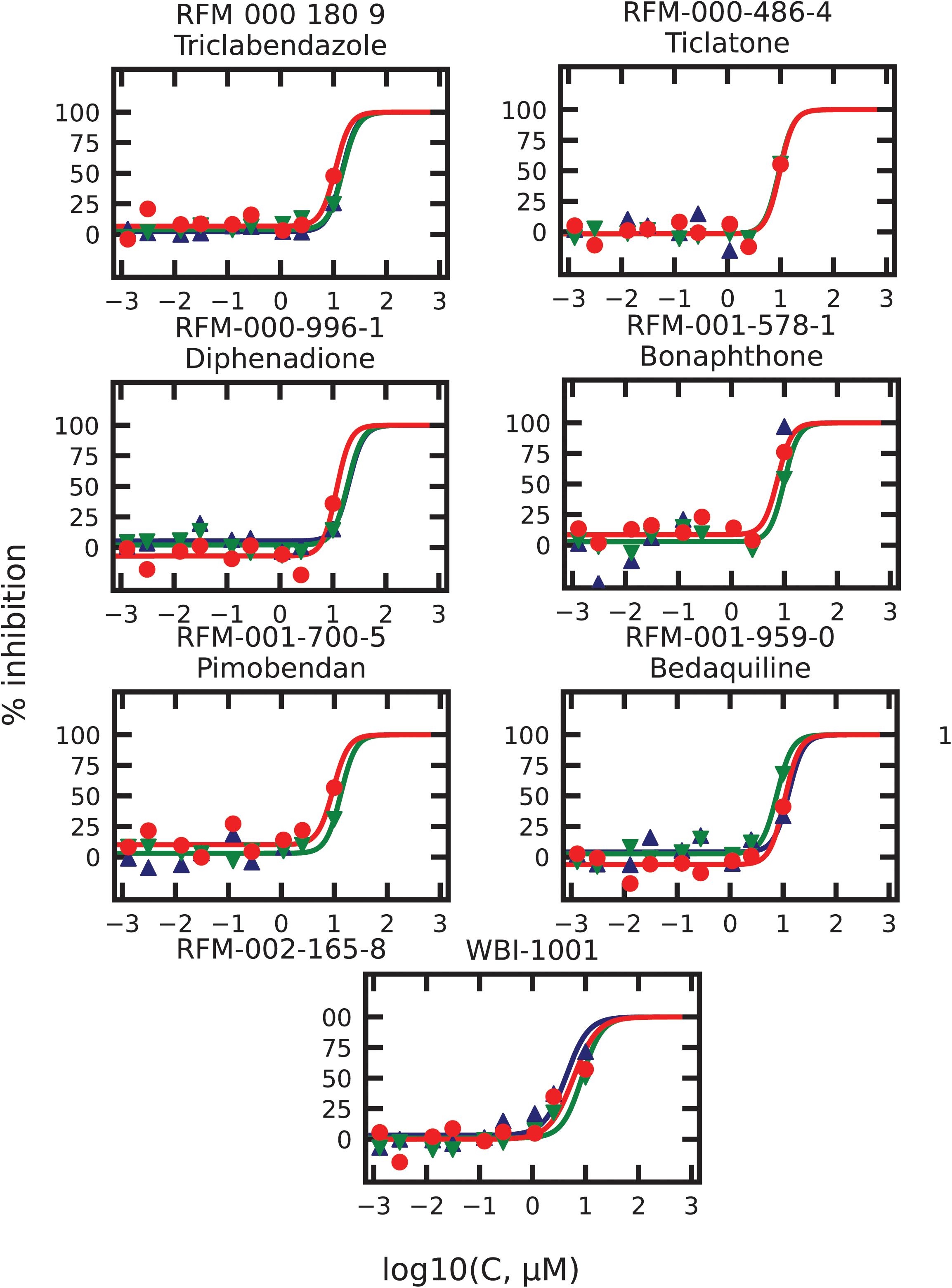
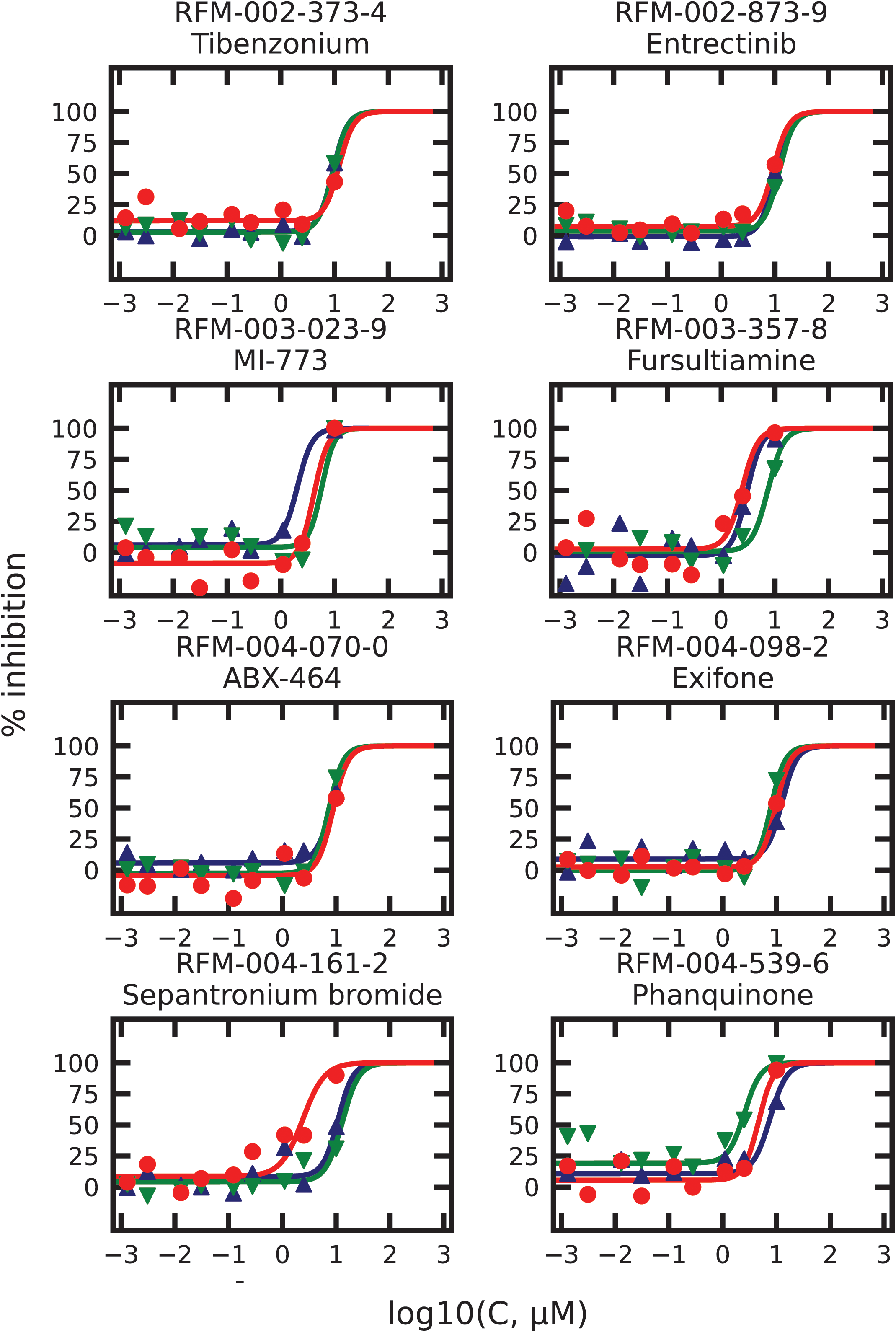
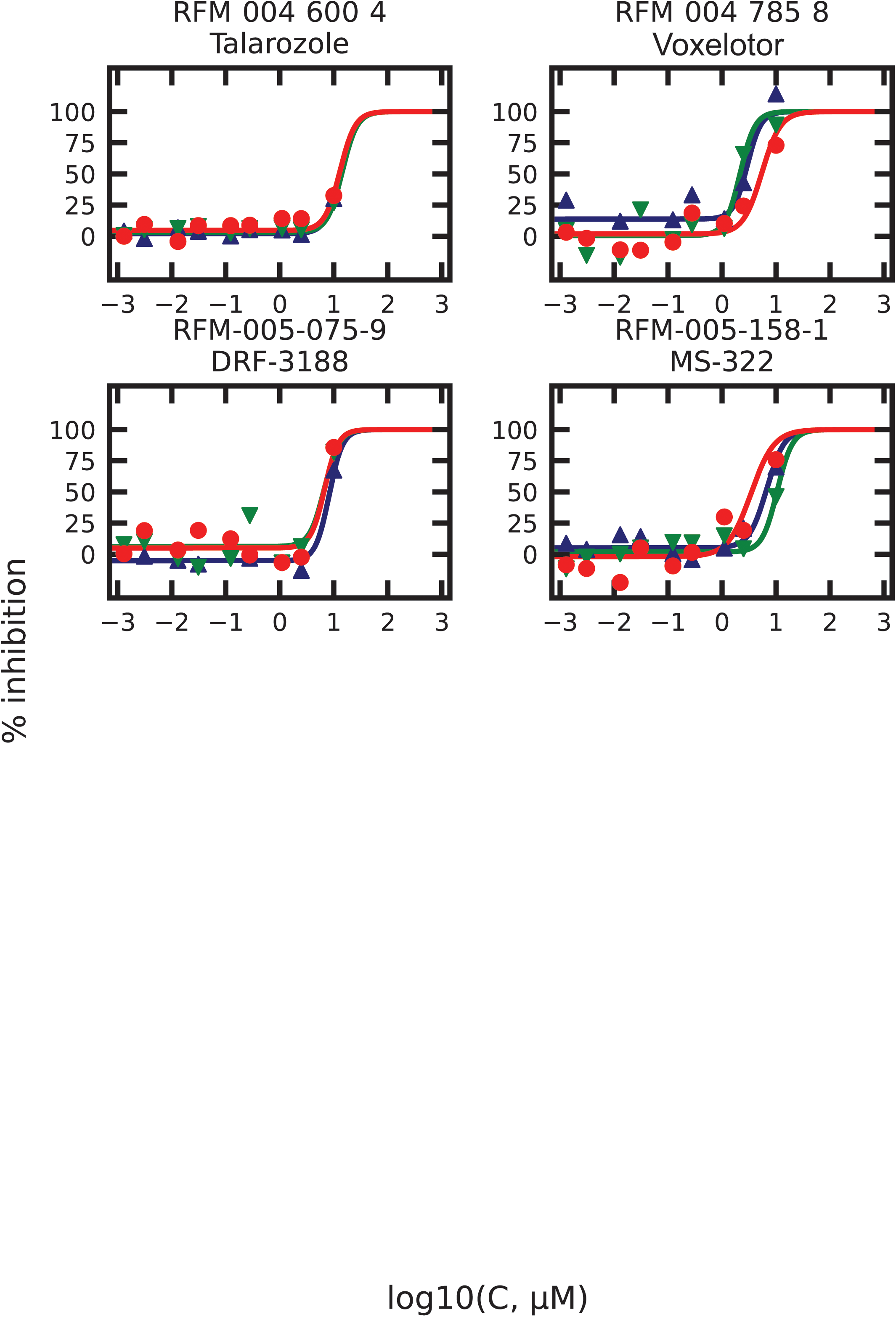
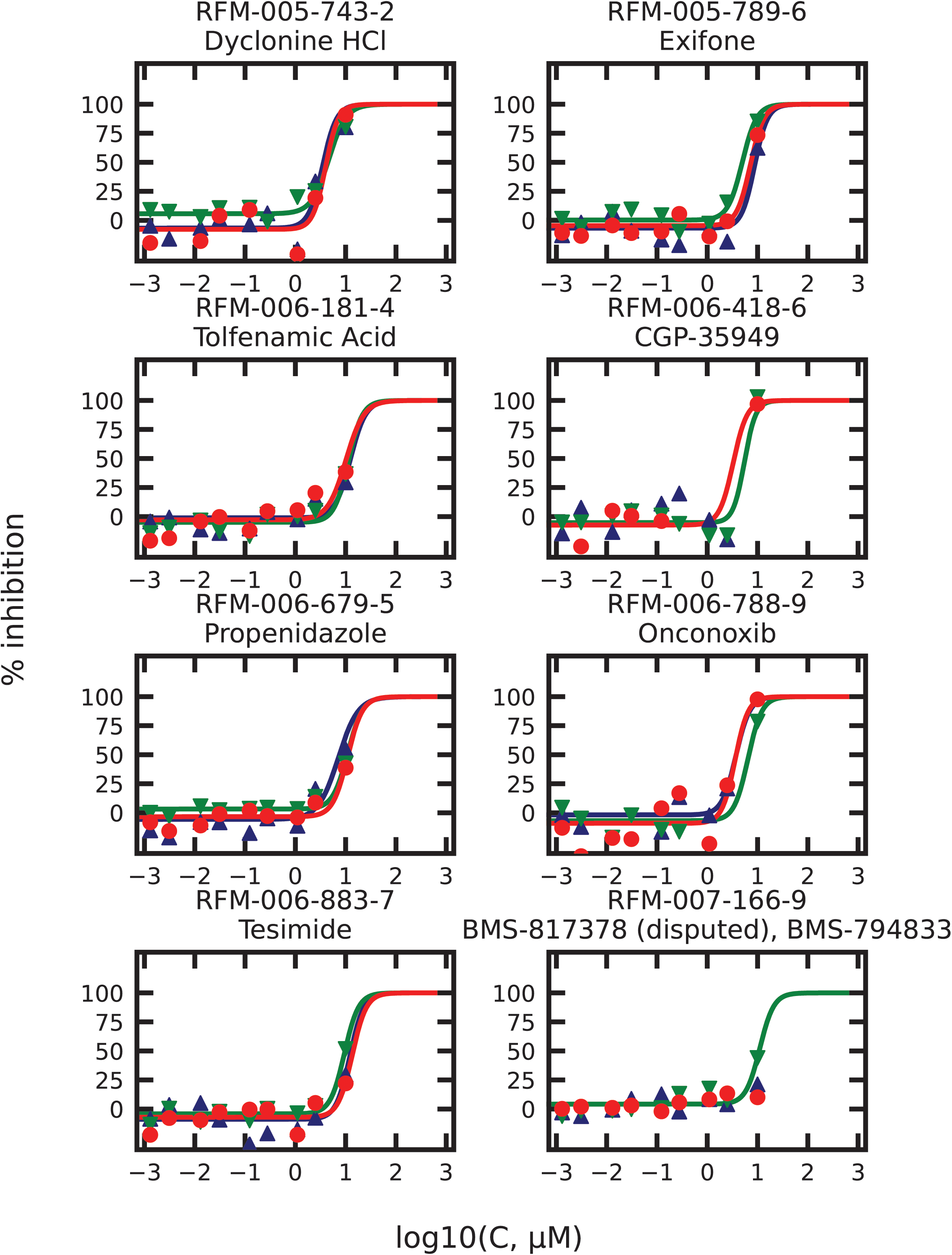
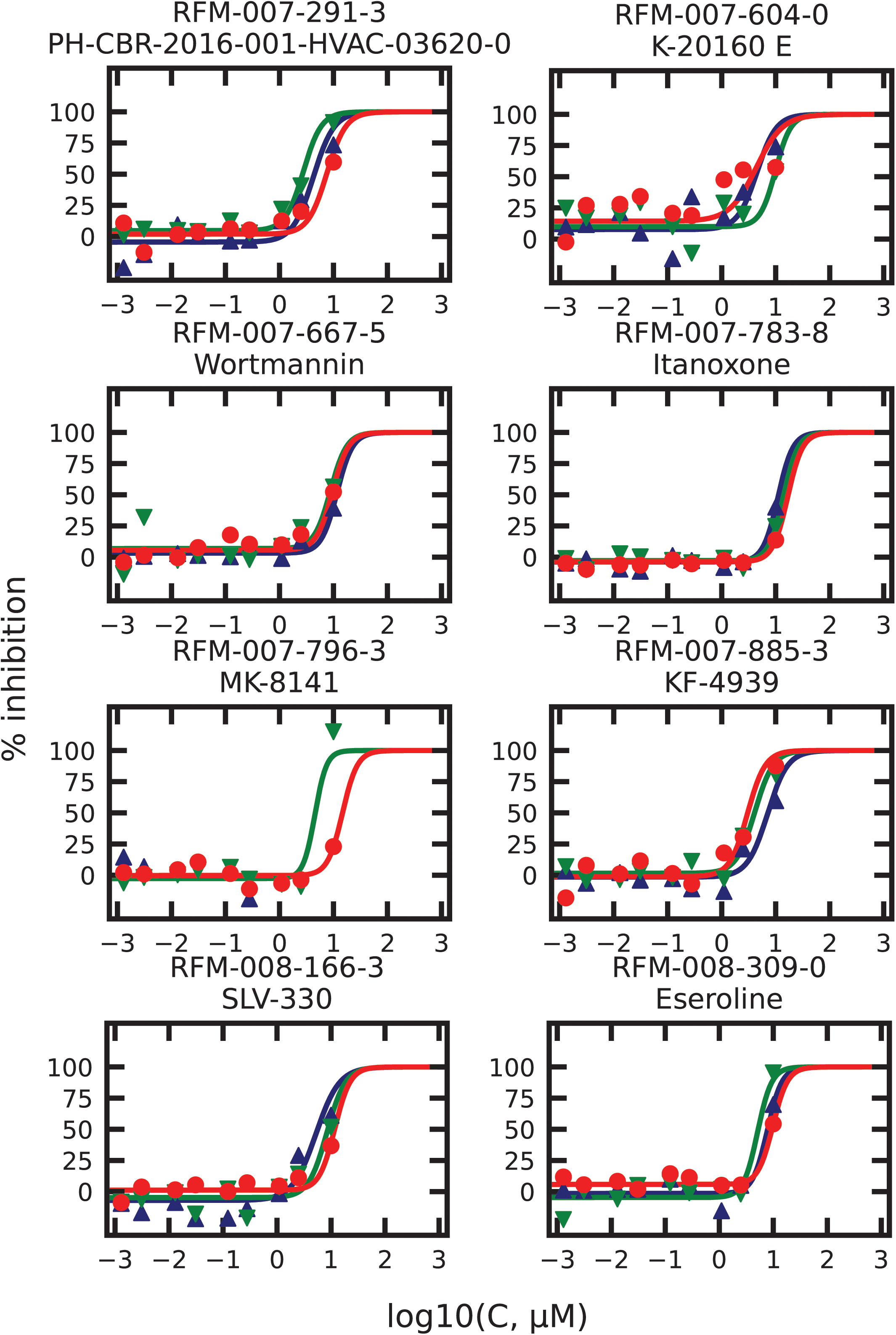
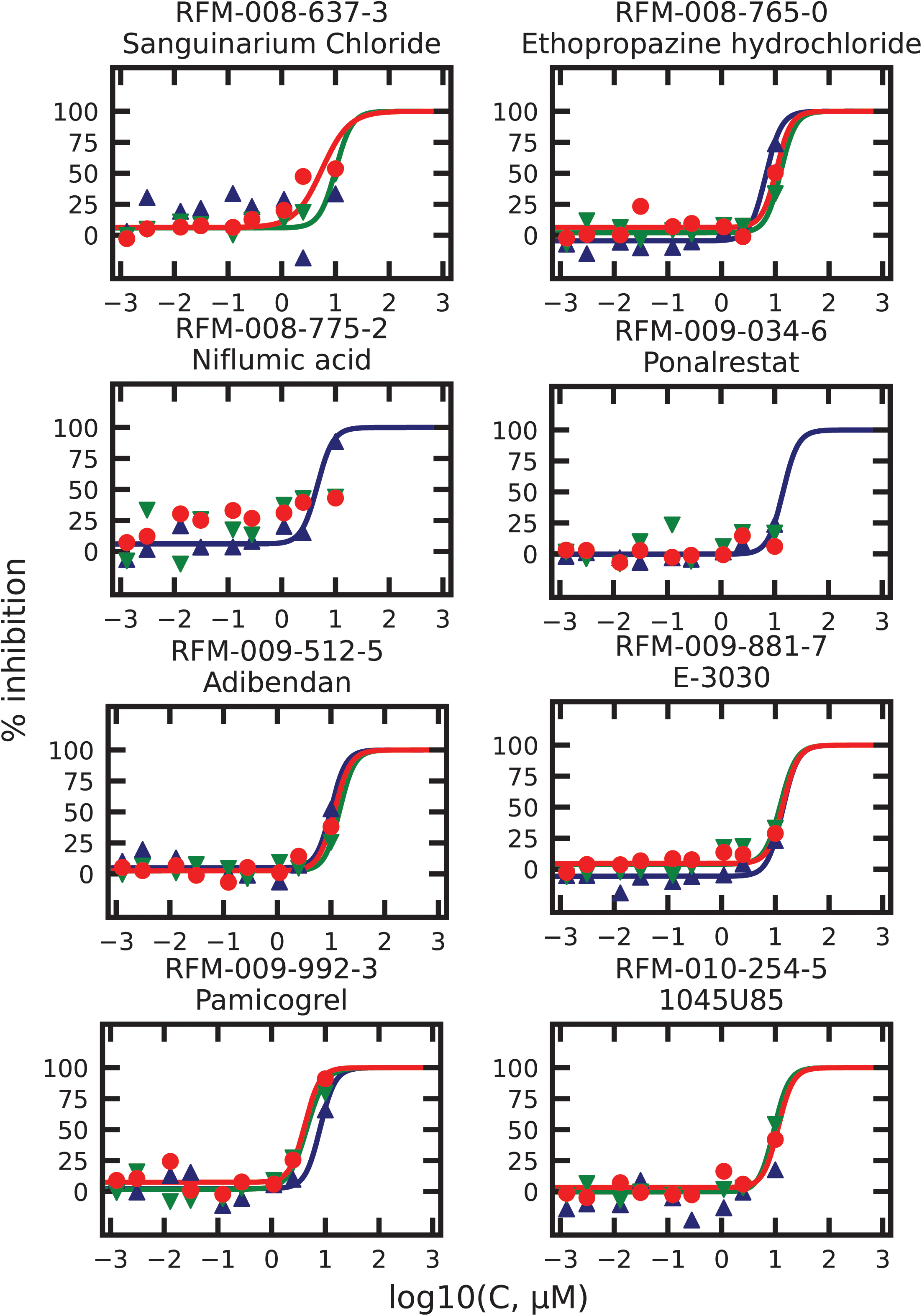
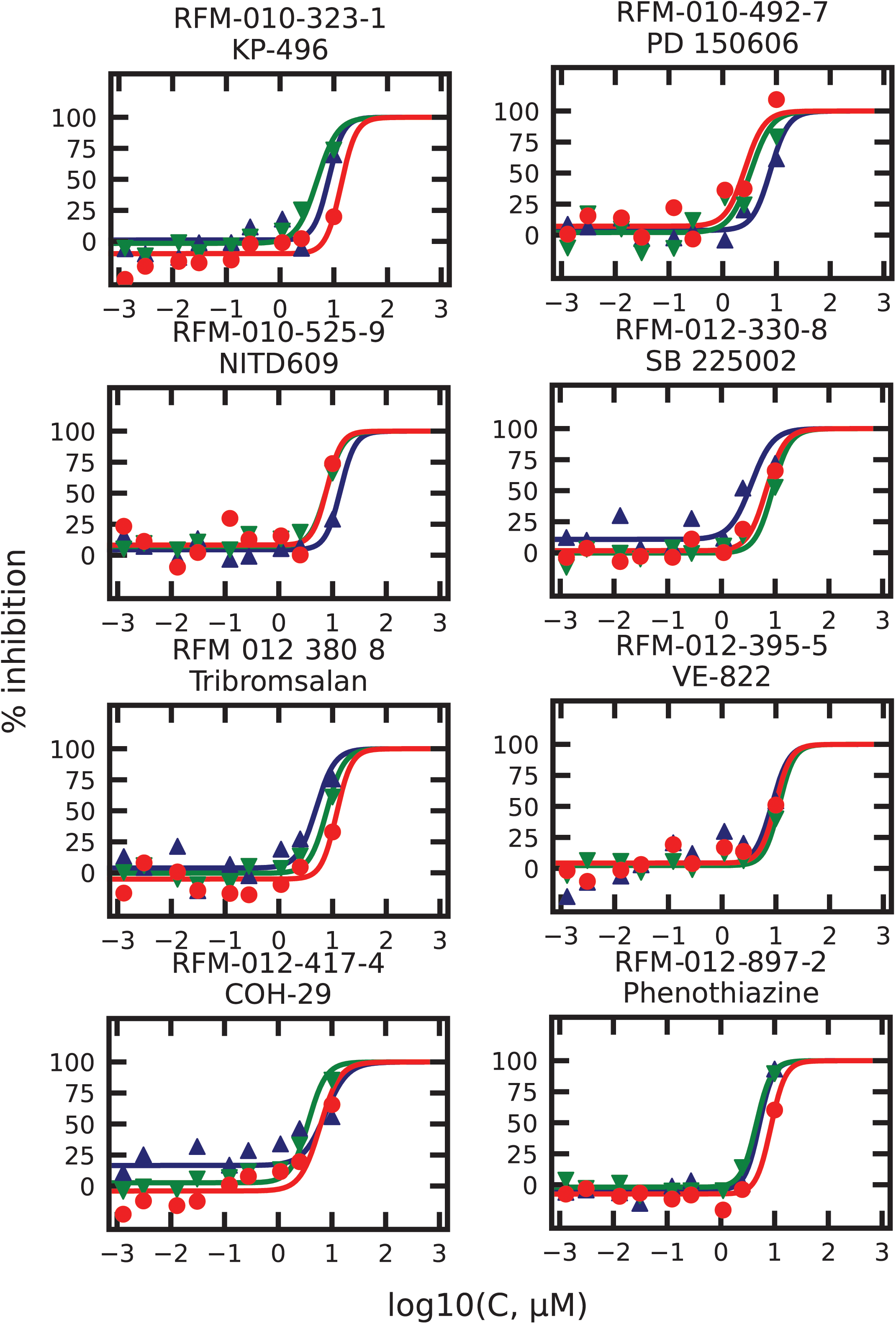

## Footnotes

* Calibr provided the compounds required for screening as spotted well plates with each well containing less than 100 nanograms. Our next step is to acquire milligram quantities of all 106 anti-sickling compounds to ascertain the detailed mechanism of inhibition for each one in order to determine which ones could be administered together as non-competitive inhibitors because they have different targets. The class of mechanism to which a particular compound belongs will be determined from measurements of cell volumes for assessing changes in intracellular HbS concentration, measurement of oxygen dissociation curves to detect preferential binding to the R quaternary conformation, and application of our laser CO photolysis assay (14) or the Wood-Sachs FRET and dynamic light scattering methods(50) on dilute HbS solutions to detect destabilization of the fiber.

† After this work was completed, we became aware of another high throughput and extremely interesting assay based on detecting pre-nuclear oligomers in dilute solutions of purified HbS using fluorescence D.K. Wood(50). In a screen of the 1280 LOPAC compound library they discovered three anti-polymerization compounds that do not alter oxygen affinity - chlormezanone, gabazine, and phosphoramidon disodium. At 500 μM concentration, these compounds also inhibit polymerization inside red cells as judged by a microfluidic rheological assay of whole blood at a hematocrit of 25%, a typical value in SCD. However, in the dilute solution assay, 100 μM of these compounds was required to observe inhibition, whereas only ∼2% of oral drugs are found in human serum at free concentrations of 100 μM or more. Moreover, chlormezanone was one of the compounds in the Calibr-Scripps library and showed no inhibition at 10 μM. With the current information it seems unlikely that any of the three compounds could be developed into oral drugs for sickle cell disease. However, we should emphasize that the demonstration of inhibition of pre-nuclear aggregation inhibiting intracellular HbS polymerization is a very important finding. In our future work on determining the mechanism of sickling inhibition of our 106 hit compounds, we will use the very important dynamic light scattering and FRET assays of Wood et al. to determine which ones act by blocking intermolecular contacts in the sickle fiber. Fortunately, we can use our Agilent Lionheart automated microscope system for these experiments because it was actually designed for fluorescence measurements.

‡ Mitapivat, one of the agents that demonstrated potential anti-sickling properties in the screen (Table 1), has been tested in a phase 1 study in patients with SCD (51). Mitapivat is an activator of red cell pyruvate kinase (PKR); the study established proof of concept that activating PKR is viable therapeutic approach for SCD, it improved hematologic parameters, increased oxygen affinity and reduced sickling in patients with SCD. Results from this study informed the design of an ongoing phase 2/3 trial of mitapivat in patients with SCD (RISE UP; NCT05031780), evaluating both improvements in Hb and frequency of sickle cell pain crises.

